# The symmetrical pattern of base-pair substitution rates across the chromosome in *Escherichia coli* has multiple causes

**DOI:** 10.1101/523696

**Authors:** Brittany A. Niccum, Heewook Lee, Wazim MohammedIsmail, Haixu Tang, Patricia L. Foster

## Abstract

Mutation accumulation experiments followed by whole-genome sequencing have revealed that for several bacterial species the rate of base-pair substitutions is not constant across the chromosome but varies in a wave-like pattern symmetrical about the origin of replication. The experiments reported here demonstrate that in *Escherichia coli* several interacting factors determine the wave. Perturbing replication timing, progression, or the structure of the terminus disrupts the pattern. Biases in error-correction by proofreading and mismatch repair are major factors. The activities of the nucleoid binding proteins, HU and Fis, are important, suggesting that mutation rates increase when highly structured DNA is replicated. These factors should apply to most bacterial, and possibly eukaryotic, genomes, and imply that different areas of the genome evolve at different rates.

**Author Summary:** In several species of bacteria the rate of single base-pair mutations is not constant across the genome, but varies in a wave-like pattern that is symmetrical about the origin of replication. Using Escherichia coli as our model system, we show that this wave is determined by the timing and progression of replication and the structure of the region where replication terminates. In addition, biases in error correction and the three dimensional structure of the DNA also are important. These factors should apply to most bacterial, and possibly eukaryotic, genomes, and imply that different areas of the genome evolve at different rates.

## Introduction

Recent studies of mutations accumulated non-selectively across bacterial chromosomes have revealed that rates of base-pair substitutions (BPSs) vary 2-to 4-fold in a wave like pattern that is mirrored in the two independently replicating halves of the chromosome. Such symmetrical patterns have been observed in mismatch-repair (MMR) defective strains of *Escherichia coli* (1), *Vibrio fischeri, V. cholerae* (2–4), *Pseudomonas fluorescens* (5), and *P. aeruginosa* (6). Such variation in mutation rates may affect the rate at which genes in different regions of the chromosome evolve, and may exert selective pressure on gene placement. Yet the causes of this variation are not known.

The fidelity of DNA replication, which, in *E. coli*, is about 1 mistake in 1000 generations (7), is determined by the intrinsic accuracy of the DNA polymerase and error correction by proofreading and mismatch repair (reviewed in (8, 9). In *E. coli*, proofreading is performed by epsilon, a subunit of the DNA polymerase III holoenzyme. If the polymerase inserts the incorrect base, epsilon’s 5’ to 3’ exonuclease activity degrades a few bases of the new strand and polymerase then re-synthesizes it. The accuracy of DNA synthesis is improved about 4000-fold by proofreading (10). Mismatch repair is performed by three proteins, MutS, MutL, and MutH. MutS recognizes a mismatch and recruits MutL. Together they find a nearby GATC site in which, in *E. coli*, the A is methylated by the Dam methylase. Because methylation lags behind replication, unmethylated As identify the “new”, and presumably error-containing, DNA strand. MutH is recruited by MutSL to the GATC site and activated to nick the unmethylated DNA strand, which is then degraded past the mismatch by the concerted activity of the UvrD helicase and one of four exonucleases. Pol III then re-synthesizes the strand. MMR improves the accuracy of DNA replication 100 to 200 fold (11).

In our previous study of the BPS density pattern in MMR-defective *E. coli* (1), we correlated mutation rates to the chromosomal sites that are affected by two nucleoid-associated proteins (NAPs), HU and Fis. We suggested that when the replication fork encounters regions of the chromosome with high superhelical density due to the binding of these NAPs, the mutation rate increases. An alternative explanation ties mutation rates to replication timing (2, 3). An intriguing hypothesis is that mutation rates vary in concert with fluctuations in dNTP concentration when the replication origin fires repeatedly during rapid growth (2).

In the work presented here, we investigate further the causes of the wave-like pattern of BPS rates. MMR-defective strains of *E. coli* additionally defective for other activities were used in mutation accumulation (MA) experiments and the mutations identified by whole genome sequencing (WGS). We also investigated the effects of different growth conditions on BPS rates. We conclude that the BPS density pattern does not have a single cause, but is the result of several factors affecting DNA replication, repair, and chromosome structure. In addition, we report that a MMR-defective *Bacillus subtilis* also has a wave-like BPS density pattern that is symmetrical about the origin of replication.

## Results

### The base-pair substitution (BPS) density pattern in mismatch repair-defective strains

In a previous paper (1) we demonstrated that the density of BPS that accumulated across the chromosome in a MMR-defective strain during an MA experiment fell into a wave-like pattern that was symmetrical about the origin. We have since performed nine additional MA experiments with MMR mutant strains, each experiment resulting in the accumulation of over 1000 BPSs, for a total of 30,061 BPSs (11). As shown in Figure 1A, the wave-like BPS density pattern was reproducible among the 10 experiments. Also as shown in Figure 1A, the mutational density pattern does not exactly match the non-interacting chromosomal macrodomains (MCs) as defined by Valens et al, 2004 (12). In particular, the wave pattern of BPSs is symmetrical about the origin whereas the Ori MC is not.

**Figure 1.**
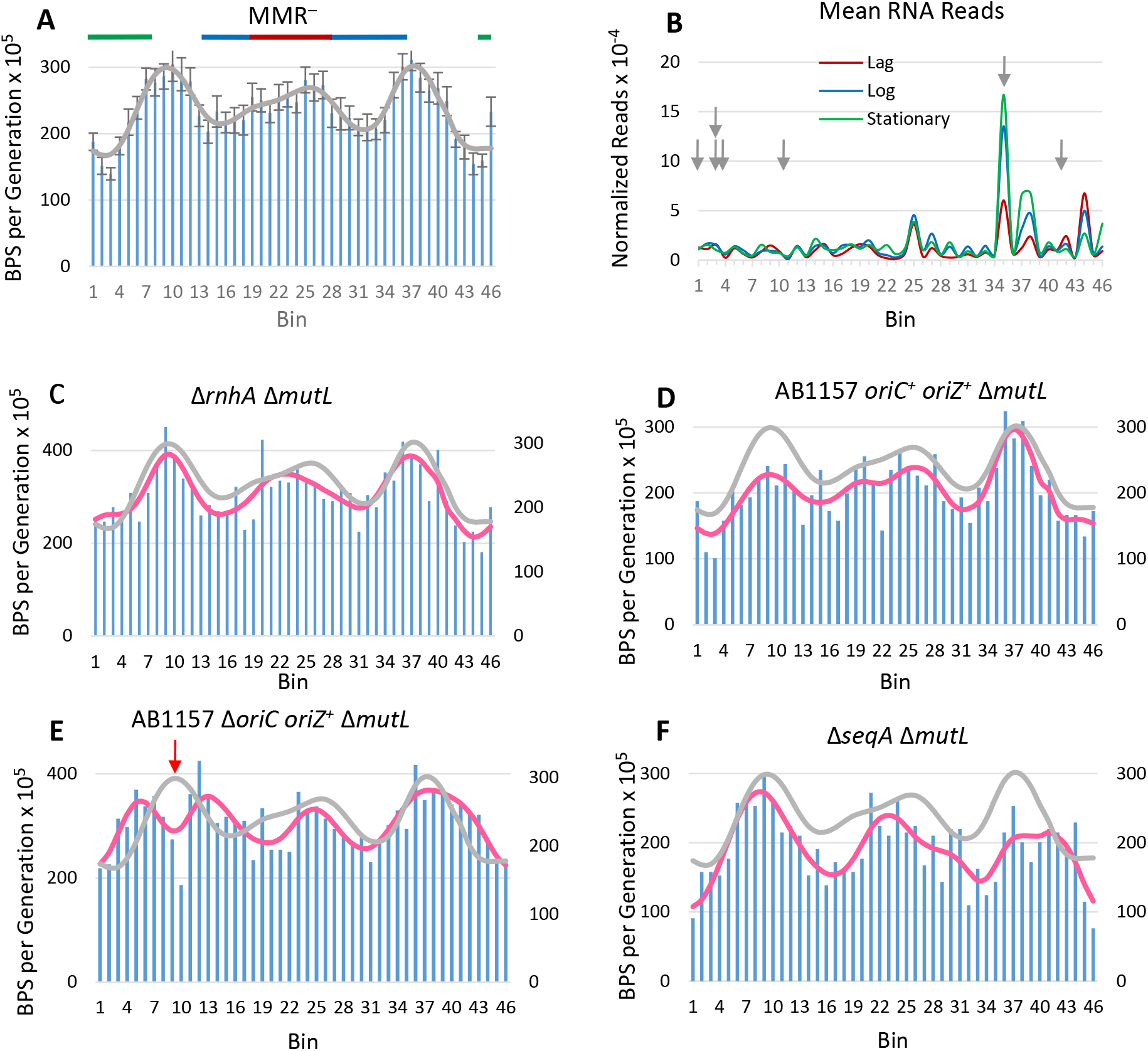
The BPS density pattern is not due to transcription but is affected by moving the origin of replication. For each plot the data were collected into 100 Kb bins starting at the origin of replication on the left and continuing clockwise around the chromosome back to the origin on the right. **Figure 1A.** The BPS density pattern of MMR-defective strains. Bars represent the mean mutation rate in each bin calculated from the BPSs collected from 10 experiments with MMR-defective strains. Error bars are the 95% CLs of the means. The grey line represents the Daubechies wavelet transform of the binned data. The chromosomal macrodomains (MCs) are indicated at the top of the plot: green, Ori MC; blue, Right and Left MC; red, Ter MC. **Figure 1B.** The Daubechies wavelet transform of the binned reads from RNA-Seq samples taken during lag, log, and stationary growth phases of the Δ*mutL* mutant strain, PFM144. The data used for the wavelet transforms were the means of three biological replicates. Arrows indicate the positions of the rRNA operons, but the reads from the ribosomal genes in those operons were not included in the plot. **Figure 1C, D, E, and F.** The BPS density patterns (bars) and Daubechies wavelet transforms (pink line) of the indicated mutant strains compared to the Daubechies wavelet transform of the combined MMR^−^ strains (grey line). See text for a description of the strains. In each plot the left-hand scale has been adjusted to bring the wavelets close together for comparison. The red arrow in 1E indicates the position of *oriZ*. Strains: 1C, PFM421; 1D, PFM430/431; 1E, PFM426; 1F, PFM533/534.

Following the analysis of Dillon et al, 2018 (2), we computed the wavelet coherence of the collected MMR^−^ data taken in the clockwise and the counterclockwise directions around the chromosome; supplementary Figure S1A shows that, except for some asymmetry at the midpoint, the wave is symmetrical across the chromosome. The coherence is greatest between 800 to 1,600 Kb per cycle (= 8 to 16 bins per cycle), which is a similar result to that found by Dillon et al, 2018 (2). We also computed the wavelet coherence of the collected MMR^−^ data against each of the experimental results discussed below, and these graphs are also given in the supplementary figures In the analyses below, the combined data from the 10 experiments with MMR-defective strains are used as the standard to which the results obtained in other genetic backgrounds are compared.

The spectrum of BPSs in the MMR-defective strains is dominated by A:T transitions at 5’NAC3’/3’NTG5’ sites (11). To test if the mutational density pattern was simply due to the distribution of these sites, we removed all the A:T transitions at 5’NAC3’/3’NTG5’ sites from the data set. As shown in supplementary Figures S5A, S6A, and Tables 1 and 2, although the mutation rate was reduced by 60% when these BPSs were removed, the remaining 12,542 BPSs fell into the same wave pattern. Thus, the variation in BPS rates must be reflective of regions of the chromosome and not the distribution of the hotspot DNA sequences.

**Table 1.** Pearson’s correlation, ρ, of each strain versus MMR^−^

| Strain | Description | Whole |  |  |  |  |  |  |  |  |  |
| --- | --- | --- | --- | --- | --- | --- | --- | --- | --- | --- | --- |
|  |  | Chromosome |  | Right Replichore |  | Left Replichore |  | Origin |  | Terminus |  |
|  |  | Bins 1 to 46 |  | Bins 1 to 23 |  | Bins 46 to 24 |  | Bins 1 to 13, 46 to 34 |  | Bins 14 to 33 |  |
| | | $\rho$ | P<sup>a</sup> | $\rho$ | P<sup>a</sup> | $\rho$ | P<sup>a</sup> | $\rho$ | P<sup>a</sup> | $\rho$ | P<sup>a</sup> |
| Collective<sup>b</sup> | MMR<sup>-</sup> | — | — | — | — | — | — | — | — | — | — |
| Collective | MMR<sup>-</sup> minus A:T ts at |  |  |  |  |  |  |  |  |  |  |
|  | NAC/NTG | 0.94 | <0.001 | 0.93 | <0.001 | 0.96 | <0.001 | 0.97 | <0.001 | 0.71 | 0.004 |
| PFM421 | $\Delta rnhA \Delta mutL$ | 0.74 | <0.001 | 0.63 | 0.002 | 0.84 | <0.001 | 0.81 | <0.001 | 0.47 | 0.075 |
| PFM669 | $\Delta mutL$ (AB1157) | 0.59 | <0.001 | 0.40 | 0.064 | 0.75 | <0.001 | 0.71 | <0.001 | 0.45 | 0.079 |
| PFM430/ 431 | *oriC<sup>+</sup> oriZ<sup>+</sup> $\Delta mutL$* |  |  |  |  |  |  |  |  |  |  |
|  | (AB1157) | 0.75 | <0.001 | 0.71 | <0.001 | 0.81 | <0.001 | 0.83 | <0.001 | 0.46 | 0.075 |
| PFM426 | $\Delta oriC oriZ+ \Delta mutL$ | | | | | | | | | | |
|  | (AB1157) | 0.39 | 0.008 | 0.21 | 0.34 | 0.62 | 0.003 | 0.40 | 0.044 | 0.41 | 0.11 |
| PFM533/ 534 | $\Delta seqA \Delta mutL$ | 0.49 | 0.001 | 0.71 | <0.001 | 0.27 | 0.23 | 0.53 | 0.006 | 0.38 | 0.14 |
| PFM5<sup>b</sup> | $\Delta mutL$ on minimal | | | | | | | | | | |
|  | medium | 0.50 | 0.001 | 0.40 | 0.069 | 0.61 | 0.003 | 0.57 | 0.003 | 0.58 | 0.029 |
| PFM343<sup>b</sup> | *ΔmutS* on minimal |  |  |  |  |  |  |  |  |  |  |
|  | medium | 0.18 | 0.22 | 0.18 | 0.41 | 0.18 | 0.41 | 0.21 | 0.31 | 0.22 | 0.38 |
| PFM343<sup>b</sup> | *ΔmutS* on diluted LB | 0.84 | <0.001 | 0.84 | <0.001 | 0.85 | <0.001 | 0.87 | <0.001 | 0.71 | 0.004 |
| PFM343<sup>b</sup> | *ΔmutS* on supplemented |  |  |  |  |  |  |  |  |  |  |
|  | minimal medium | 0.81 | <0.001 | 0.82 | <0.001 | 0.81 | <0.001 | 0.83 | <0.001 | 0.65 | 0.008 |
| PFM342<sup>b</sup> | *ΔmutS* at low |  |  |  |  |  |  |  |  |  |  |
|  | temperature | 0.57 | <0.001 | 0.60 | 0.003 | 0.53 | 0.013 | 0.69 | <0.001 | -0.02 | 0.95 |
| PFM799 | *ΔnrdR ΔmutL* | 0.49 | 0.001 | 0.58 | 0.005 | 0.39 | 0.080 | 0.69 | <0.001 | -0.31 | 0.24 |
| PFM677 | *Δrep ΔmutL* | 0.48 | 0.001 | 0.49 | 0.022 | 0.48 | 0.027 | 0.62 | 0.001 | -0.12 | 0.67 |
| PFM256 | *Δtus ΔmutL* | 0.82 | <0.001 | 0.79 | <0.001 | 0.85 | <0.001 | 0.87 | <0.001 | 0.70 | 0.004 |
| PFM257 | *ΔmatP ΔmutL* | 0.40 | 0.007 | 0.60 | 0.004 | 0.21 | 0.34 | 0.70 | <0.001 | -0.47 | 0.075 |
| PFM422 | *ΔrecA ΔmutL* | 0.71 | <0.001 | 0.70 | <0.001 | 0.73 | <0.001 | 0.85 | <0.001 | 0.06 | 0.84 |
| PFM424 | *ΔrecA ΔmutS* | 0.80 | <0.001 | 0.78 | <0.001 | 0.83 | <0.001 | 0.91 | <0.001 | 0.44 | 0.082 |
| PFM456 | *ΔrecB ΔmutL* | 0.82 | <0.001 | 0.87 | <0.001 | 0.79 | <0.001 | 0.89 | <0.001 | 0.39 | 0.12 |
| PFM118 | *ΔumuDC ΔdinB ΔmutL* | 0.59 | <0.001 | 0.62 | 0.003 | 0.55 | 0.010 | 0.61 | 0.001 | 0.69 | 0.004 |
| PFM120 | *lexA3 ΔsulA ΔmutL* | 0.82 | <0.001 | 0.82 | <0.001 | 0.83 | <0.001 | 0.83 | <0.001 | 0.80 | 0.001 |
| PFM259 | *ΔhupB ΔmutL* | 0.73 | <0.001 | 0.70 | <0.001 | 0.77 | <0.001 | 0.81 | <0.001 | 0.57 | 0.031 |
| PFM258 | *ΔhupA ΔmutL* | 0.49 | 0.001 | 0.43 | 0.053 | 0.58 | 0.005 | 0.67 | <0.001 | -0.23 | 0.38 |
| PFM317/ 318 | *Δfis ΔmutL* | 0.51 | <0.001 | 0.61 | 0.003 | 0.42 | 0.060 | 0.67 | <0.001 | -0.30 | 0.25 |
| PFM482 | *Δfis ΔmutS* | 0.49 | 0.001 | 0.31 | 0.15 | 0.62 | 0.003 | 0.65 | <0.001 | -0.28 | 0.27 |
| PFM741 | *Δhns ΔmutL* | 0.81 | <0.001 | 0.81 | <0.001 | 0.81 | <0.001 | 0.88 | <0.001 | 0.46 | 0.075 |
| PFM713 | *Δdps ΔmutL* | 0.80 | <0.001 | 0.84 | <0.001 | 0.78 | <0.001 | 0.89 | <0.001 | 0.47 | 0.075 |
| PFM163<sup>c</sup> | *mutD5* | 0.76 | <0.001 | 0.79 | <0.001 | 0.74 | <0.001 | 0.82 | <0.001 | 0.78 | 0.001 |
| PFM165/397/399<sup>c</sup> | *mutD5 ΔmutL* | 0.64 | <0.001 | 0.76 | <0.001 | 0.52 | 0.014 | 0.86 | <0.001 | -0.46 | 0.075 |
| Collective<sup>b</sup> | Wild type | 0.36 | 0.014 | 0.41 | 0.061 | 0.31 | 0.17 | 0.37 | 0.065 | 0.51 | 0.060 |
| Collective<sup>d</sup> | *Bacillus subtilis* |  |  |  |  |  |  |  |  |  |  |
|  | *mutS::Tn10* | 0.52 | <0.001 | 0.63 | 0.003 | 0.40 | 0.068 | 0.74 | <0.001 | -0.52 | 0.055 |

<sup>a</sup> P values were adjusted for multiple comparisons using the Benjamini-Hochberg method (71) with the false discovery rate set at 25%
<sup>b</sup> Data are from (11)
<sup>c</sup> Data are from (10)
<sup>d</sup> Data are combined from four MA experiments with DK2140 (NCIB3610 *mutS::Tn10*), DK2141 (NCIB3610 *mutS::Tn10*), DK2142 (NCIB3610 *mutS::Tn10 ΔcomI*), and DK2143 (NCIB3610 *mutS::Tn10 ΔcomI*). There were no differences in mutation rates or spectra among these strain

**Table 2.**
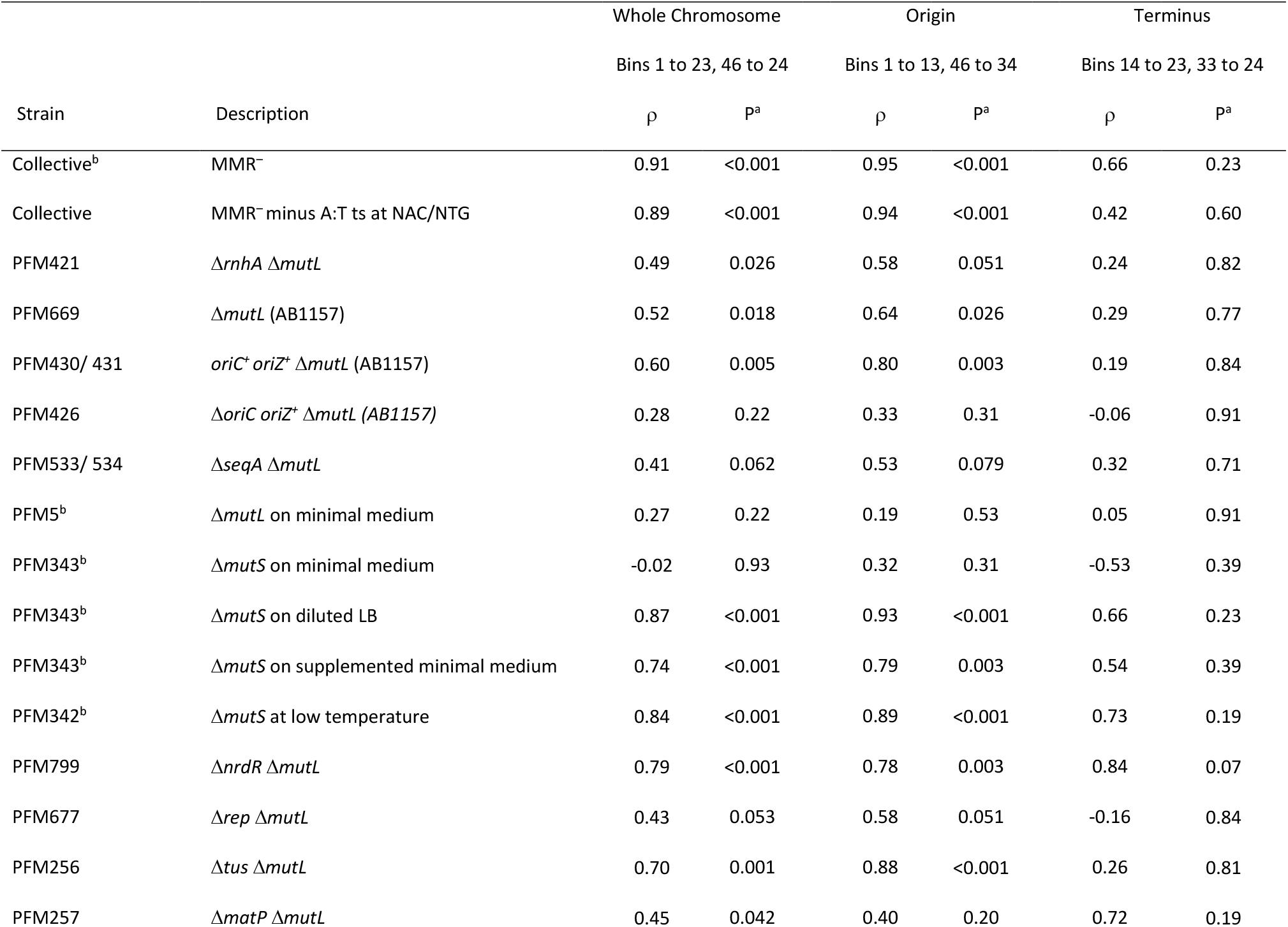
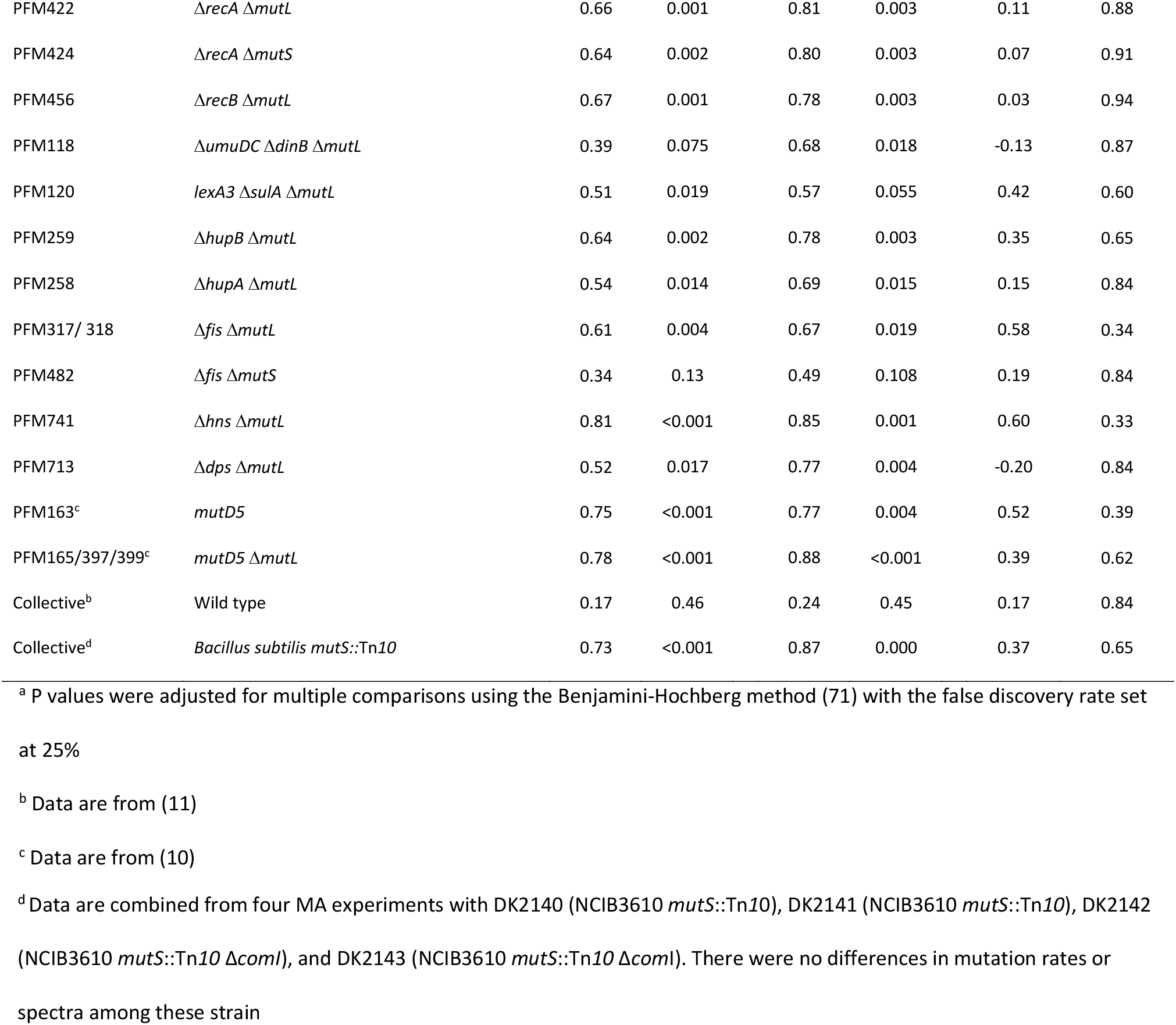
Pearson’s correlation, ρ, of right versus left replichore for each strain

| Strain | Description | Whole Chromosome |  | Origin |  | Terminus |  |
| --- | --- | --- | --- | --- | --- | --- | --- |
|  |  | Bins 1 to 23, 46 to 24 |  | Bins 1 to 13, 46 to 34 |  | Bins 14 to 23, 33 to 24 |  |
| | | $\rho$ | P<sup>a</sup> | $\rho$ | P<sup>a</sup> | $\rho$ | P<sup>a</sup> |
| Collective<sup>b</sup> | MMR<sup>-</sup> | 0.91 | <0.001 | 0.95 | <0.001 | 0.66 | 0.23 |
| Collective | MMR<sup>-</sup> minus A:T ts at NAC/NTG | 0.89 | <0.001 | 0.94 | <0.001 | 0.42 | 0.60 |
| PFM421 | $\Delta rnhA \Delta mutL$ | 0.49 | 0.026 | 0.58 | 0.051 | 0.24 | 0.82 |
| PFM669 | $\Delta mutL$ (AB1157) | 0.52 | 0.018 | 0.64 | 0.026 | 0.29 | 0.77 |
| PFM430/ 431 | $oriC^+ oriZ^+ \Delta mutL$ (AB1157) | 0.60 | 0.005 | 0.80 | 0.003 | 0.19 | 0.84 |
| PFM426 | $\Delta oriC oriZ^+ \Delta mutL$ (AB1157) | 0.28 | 0.22 | 0.33 | 0.31 | -0.06 | 0.91 |
| PFM533/ 534 | $\Delta seqA \Delta mutL$ | 0.41 | 0.062 | 0.53 | 0.079 | 0.32 | 0.71 |
| PFM5<sup>b</sup> | $\Delta mutL$ on minimal medium | 0.27 | 0.22 | 0.19 | 0.53 | 0.05 | 0.91 |
| PFM343<sup>b</sup> | $\Delta mutS$ on minimal medium | -0.02 | 0.93 | 0.32 | 0.31 | -0.53 | 0.39 |
| PFM343<sup>b</sup> | $\Delta mutS$ on diluted LB | 0.87 | <0.001 | 0.93 | <0.001 | 0.66 | 0.23 |
| PFM343<sup>b</sup> | $\Delta mutS$ on supplemented minimal medium | 0.74 | <0.001 | 0.79 | 0.003 | 0.54 | 0.39 |
| PFM342<sup>b</sup> | $\Delta mutS$ at low temperature | 0.84 | <0.001 | 0.89 | <0.001 | 0.73 | 0.19 |
| PFM799 | $\Delta nrdR \Delta mutL$ | 0.79 | <0.001 | 0.78 | 0.003 | 0.84 | 0.07 |
| PFM677 | $\Delta rep \Delta mutL$ | 0.43 | 0.053 | 0.58 | 0.051 | -0.16 | 0.84 |
| PFM256 | $\Delta tus \Delta mutL$ | 0.70 | 0.001 | 0.88 | <0.001 | 0.26 | 0.81 |
| PFM257 | $\Delta matP \Delta mutL$ | 0.45 | 0.042 | 0.40 | 0.20 | 0.72 | 0.19 |
| PFM422 | *ΔrecA ΔmutL* | 0.66 | 0.001 | 0.81 | 0.003 | 0.11 | 0.88 |
| PFM424 | *ΔrecA ΔmutS* | 0.64 | 0.002 | 0.80 | 0.003 | 0.07 | 0.91 |
| PFM456 | *ΔrecB ΔmutL* | 0.67 | 0.001 | 0.78 | 0.003 | 0.03 | 0.94 |
| PFM118 | *ΔumuDC ΔdinB ΔmutL* | 0.39 | 0.075 | 0.68 | 0.018 | -0.13 | 0.87 |
| PFM120 | *lexA3 ΔsulA ΔmutL* | 0.51 | 0.019 | 0.57 | 0.055 | 0.42 | 0.60 |
| PFM259 | *ΔhupB ΔmutL* | 0.64 | 0.002 | 0.78 | 0.003 | 0.35 | 0.65 |
| PFM258 | *ΔhupA ΔmutL* | 0.54 | 0.014 | 0.69 | 0.015 | 0.15 | 0.84 |
| PFM317/ 318 | *Δfis ΔmutL* | 0.61 | 0.004 | 0.67 | 0.019 | 0.58 | 0.34 |
| PFM482 | *Δfis ΔmutS* | 0.34 | 0.13 | 0.49 | 0.108 | 0.19 | 0.84 |
| PFM741 | *Δhns ΔmutL* | 0.81 | <0.001 | 0.85 | 0.001 | 0.60 | 0.33 |
| PFM713 | *Δdps ΔmutL* | 0.52 | 0.017 | 0.77 | 0.004 | -0.20 | 0.84 |
| PFM163<sup>c</sup> | *mutD5* | 0.75 | <0.001 | 0.77 | 0.004 | 0.52 | 0.39 |
| PFM165/397/399<sup>c</sup> | *mutD5 ΔmutL* | 0.78 | <0.001 | 0.88 | <0.001 | 0.39 | 0.62 |
| Collective<sup>b</sup> | Wild type | 0.17 | 0.46 | 0.24 | 0.45 | 0.17 | 0.84 |
| Collective<sup>d</sup> | *Bacillus subtilis mutS::Tn10* | 0.73 | <0.001 | 0.87 | 0.000 | 0.37 | 0.65 |

<sup>a</sup> P values were adjusted for multiple comparisons using the Benjamini-Hochberg method (71) with the false discovery rate set at 25%
<sup>b</sup> Data are from (11)
<sup>c</sup> Data are from (10)
<sup>d</sup> Data are combined from four MA experiments with DK2140 (NCIB3610 *mutS*::Tn10), DK2141 (NCIB3610 *mutS*::Tn10), DK2142 (NCIB3610 *mutS*::Tn10 $\Delta$ *comI*), and DK2143 (NCIB3610 *mutS*::Tn10 $\Delta$ *comI*). There were no differences in mutation rates or spectra among these strain

### Transcription

One obvious hypothesis is that the mutational density pattern reflects transcriptional patterns. Mutation rates have been reported to be both increased (13) and decreased (14) by high levels of transcription. In a previous paper (7) we reported that highly expressed genes had normal mutation rates, a finding that was confirmed in a recent study using deep sequencing to detect mutations (15). To determine if, nonetheless, transcription influences the wave pattern, we used RNA-Seq to quantitate the RNA levels in a Δ*mutL* mutant strain in the lag, exponential, and stationary phases of growth. These results should be representative of the cells in colonies during our MA experiments. We then compared these results to the BPS density pattern by binning the RNA-Seq results into the same bins as used for mutational data. As shown in Figure 1B and supplementary Figure S1B, there was no similarity in the two data sets.

Because ribosomal operons are homologous, we could not call SNPs in these genes. The RNA reads from the genes in ribosomal operons were also removed from the RNA-Seq data, but their positions have been indicated in Figure 1B. Interestingly, bin 35, which includes the ribosomal *rrnG* operon, has a high number of RNA reads even in the absence of reads from the *rrnG* genes. This high level of expression is almost exclusively due to the *ssrA* gene, which encodes transfer-messenger RNA (16), that is highly expressed under all three conditions (the RNA-Seq data will be further analyzed in a subsequent paper). The BPS density pattern of several of the strains that will be discussed below tends to reach a minimum in this general area, but it is as often at bin 33 as at bin 35. Given that the bin size is 100 Kb, it seems unlikely that the high level of transcription level of *ssrA*, located in the middle of bin 35, is causing these patterns.

### Replication Initiation

The BPS density pattern is centered on *oriC*, the origin of replication, and over many experiments in different genetic backgrounds the pattern around the origin has proved to be stable. In both replichores BPS rates decline to a minimum about 300 Kb from the origin and then increase until a peak is reached about 900 Kb from the origin. Mutation rates then fall again and reach a minimum about 3/5^th^ of the distance along each replichore.

We tested whether replication initiation was responsible for the maintenance of this pattern by performing MA experiments with strains with errant replication start sites. The *rnhA* gene encodes RNase H1, which degrades RNA-DNA hybrids; in the absence of RNase H1, persistent R-loops can initiate aberrant DNA replication and disrupt normal fork migration (17, 18). But an MA experiment with a Δ*rnhA* Δ*mutL* mutant strain showed no difference in the BPS density pattern from that of the Δ*mutL* mutant strain (Figure 1C, Supplementary Figure S1C, Tables 1 and 2), indicating that aberrant replication Initiation does not influence where mutations occur, at least not when a powerful *oriC* is present.

To further test the influence of replication initiation on the mutational density pattern, we performed MA experiments with strains that have a 5.1 Kb region containing *oriC* moved to the midpoint of the right replichore, where it is called *oriZ* (19). These strains are derived from *E. coli* K12 strain AB1157, instead of MG1655 strain, the ancestor of our MA strains, and have a large inversion in the right replichore that relieves the head-on collision between replication initiating at *oriZ* and transcription of the *rrnCABE* operon (20). As a control we created an AB1157 Δ*mutL* mutant strain, which had the similar wave-like BPS density pattern as our MMR^−^ strains, indicating that the pattern is intrinsic to *E. coli* K12, not specific to sub-strain MG1655 (Supplementary Figures S5B, S6B, Tables 1 and 2). We performed MA experiments on Δ*mutL* derivatives of two additional strains: one with both *oriZ* and *oriC* (WX320 Δ*mutL*), and one with only *oriZ* (WX340 Δ*mutL*). The strain that contained two origins had a similar BPS density pattern as the MMR^−^ strains (Figure 1D, Supplementary Figure S1D, Tables 1 and 2), suggesting that, under our conditions, firing of *oriZ* could not overcome the influence of *oriC*. However, the strain containing only *oriZ* showed a decrease in BPS rate in the 200Kb-area surrounding the new origin, similar to that normally observed about *oriC* (Figure 1E, Supplementary Figure S1E, Tables 1 and 2).

### SeqA

As mentioned above, in *E. coli* the adenines in GATC sites are methylated by the Dam methylase. The SeqA protein binds to hemimethylated GATC sites, many of which are clustered around OriC, and, by so doing, SeqA occludes the replication initiation protein, DnaA, hindering origin firing. Sequestering of the origin persists for about one third of a generation; however, the mechanism of relief is not clear. In the absence of SeqA, unregulated initiation presumably results in over-replication, at least when cells are rapidly growing in rich medium (21). Downstream events, such as replication fork collapse, add to the phenotypes of *seqA* mutant cells (22).

Loss of SeqA affects chromosomal structure in areas distant from OriC. By binding to hemimethylated DNA, SeqA forms complexes behind the replication fork as it progresses around the chromosome (21). In addition, SeqA binds to areas of the chromosome with closely spaced GATC sites, as well as to particular genes regulated by GATC methylation (23). In the absence of SeqA the superhelicity of the chromosome increases, the nucleoid condenses (24), and transcription is altered (25).

To test whether SeqA affects the BPS density pattern, we performed an MA experiment with a Δ*mutL* Δ*seqA* mutant strain. As shown in Figure 1F, Supplementary Figure S1F, and Tables 1 and 2, loss of SeqA somewhat amplified the BPS density pattern of the right replichore, but the pattern was still highly correlated to that of right replichore of the MMR-defective strains. However, the pattern in the left replichore was disrupted in the Δ*seqA* Δ*mutL* mutant strain. Based on chromatin immunoprecipitation analysis, this area of the left replichore is not targeted by SeqA to a greater extent than the same area of the right replichore (23, 26), suggesting that the disruption of the mutational pattern is not due to loss of binding by SeqA. As shown in Figure 1B, bins 44, 38, 37, and 35 contain a number of highly expressed genes; in addition, bins 42 and 35 contain highly transcribed ribosomal RNA genes. Thus, we hypothesize that loss of SeqA makes the replication machinery particularly susceptible to interference by transcription, disrupting the mutational pattern. If this hypothesis is correct, the interference apparently makes replication more accurate, perhaps by slowing the speed of DNA polymerase.

### Replication Fork Progression

In a recent report, Dillon *et al*., 2018 (2), hypothesized that the periodic variation in mutation rates across the chromosome was tied to the timing of DNA replication. Cellular levels of dNTPs are controlled by ribonucleotide reductase (RNR), the expression of which increases when the origin fires (27, 28) }. In fast-growing bacteria in which new rounds of replication are initiated before cell division, the levels of dNTPs should be high when each origin fires, but then fall as progression of the multiple forks dilute the dNTPs. High levels of dNTPs are predicted to increase the probability of misincorporation, and thus the mutation rate, whereas low levels of dNTPs should slow replication and improve fidelity. This systematic fluctuation in dNTP levels could account for the pattern of mutational density across the chromosome (2).

This hypothesis can be tested by examining the wave pattern from several experiments that we have already published (11). The number of replicating chromosomes is a positive function of the cell’s growth rate (29). When Δ*mutL* or Δ*mutS* mutant strains were grown on glucose minimal medium, which reduces the growth rate about 2-fold relative to growth on LB, the BPS density pattern became chaotic (Figure 2A, Supplementary Figures S2A, S5C, S6C, Tables 1 and 2). Supplementing the minimal medium with just enough LB to increase the growth rate to normal restored the wave-like BPS density pattern (Figure 2B, Supplementary Figure S2B, Tables 1 and 2). Growing the cells on diluted LB, on which the growth rate was the same as on minimal glucose medium, also preserved the wave-like BPS density pattern (Figure 2C, Supplementary Figure S2C, Tables 1 and 2). When the cells were grown at a lower temperature, which also reduced the growth rate 2-fold, the overall shape of the BPS density pattern was retained, but the increases in BPS rates that normally peak at about 900 Kb from each side of the origin were shifted about 200 Kb further from OriC (Figure 2D, Supplementary Figure S2D, Tables 1 and 2). In addition, the magnitude of the fluctuation of the BPS rate across the chromosome was doubled. Thus, it appears that growth rate *per se* is not a major determinant of the BPS density pattern, but other factors, such as the composition of the growth medium, are also important. For example, the expressions and DNA binding characteristics of nucleoid associated proteins (NAPs) are different under different growth conditions (30–32)

**Figure 2.**
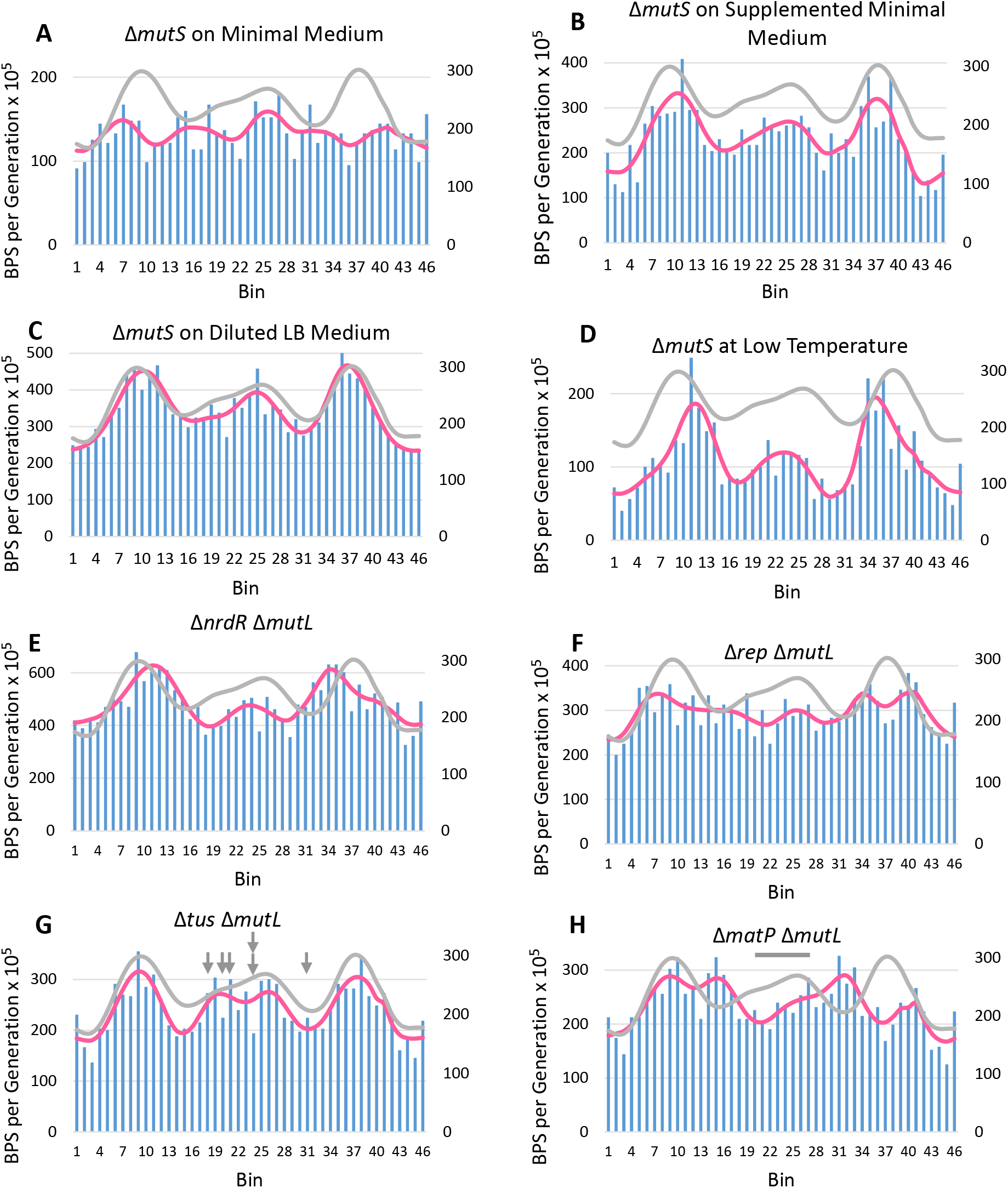
Altering the progression of DNA replication affects the BPS density pattern. For each plot the data were collected into 100 Kb bins starting at the origin of replication on the left and continuing clockwise around the chromosome back to the origin on the right. The BPS density patterns (bars) and Daubechies wavelet transforms (pink lines) of the indicated strains are compared to the Daubechies wavelet transform of the combined MMR^−^ strains (grey line). In each plot the left-hand scale has been adjusted to bring the wavelets close together for comparison. **Figure 2A, B, C, and D.** Altering the cellular growth rate by growing a Δ*mutS* mutant strain on different medium and at different temperatures differentially affects the BPS density pattern. In the experiments shown in Plot A, C, and D, the cells grew at about half the normal rate, whereas in the experiment shown in plot B they grew at the normal rate. Strains: 2A, 2B, and 2C, PFM343; 2D, PFM342. **Figure 2E and F.** Dysregulation of dNTP levels and loss of Rep, the auxiliary replication helicase, affect the BPS density pattern. 2E, PFM799; 2F, PFM677. **Figure 2G and H.** Loss of the Tus antihelicase has no effect, but loss of the terminus organizing protein, MatP, changes the BPS density pattern across the chromosome. The arrows in Figure 2G mark the major Ter sites where Tus binds. The bar in Figure 2H shows the region in which MatP binds. Strains: 2G, PFM256; 2H, PFM257.

In *E. coli* growing aerobically dNTP levels are regulated by a Class Ia RNR encoded by the *nrdA* and *nrdB* genes. Although most of this regulation appears to be due to DnaA interactions (28), the NrdR repressor, which regulates a poorly expressed Class Ib RNR, also regulates *nrdAB* transcription; loss of NrdR results in increased expression of RNR throughout the cell cycle (33). Increased RNR should result in increased dNTP levels, and, indeed, the Δ*nrdR* Δ*mutL* mutant strain had twice the mutation rate as the MMR^−^ strains, as expected when dNTP levels are high (Supplementary Table S1). As shown in Figure 2E, Supplementary Figure S2E, Tables 1 and 2, loss of NrdR did not change the basic BPS density pattern, but the peak rate was shifted away from the origin about 200 Kb on each side. This pattern was similar to that observed when cells were grown at low temperature, as described above.

Replication fork progression is aided by the accessory replication helicase, Rep, and, in its absence, the time required for chromosome duplication is doubled (34). Rep removes proteins bound to the DNA in front of the fork (34, 35). While these nucleoproteins are primarily transcription complexes (36, 37), Rep could also free the DNA of blocking NAPs. In addition, Rep aids in restarting replication forks after they stall or collapse (38, 39). As shown in Figure 2F, Supplementary Figure S2F, Tables 1 and 2, with the exception of a region close to the origin, loss of Rep disrupted the BPS density pattern across the chromosome, suggesting that slowing or stalling the fork results in a distribution of BPS that is essentially random.

### Replication Termination

The results from almost all the strains tested show an increase in the BPS rate in the region where replication terminates. The pattern of this increase varies somewhat among experiments. Usually there are two unequal peaks, as shown in Figure1A, but in some experiments these peaks are better defined and of equal heights and occasionally there is just one peak. We do not know the source of this variation, but suspect it is simply random noise.

Replication terminates approximately 180° from the origin in a 1200 Kb region bounded by replication pause (Ter) sites; this region extends from bin 18 to bin 31 in our figures. The anti-helicase Tus protein binds to the Ter sites and allows each replication fork to enter but not to exit, creating a replication fork “trap”, within which the two forks fuse and the chromosome dimer is resolved (40). To determine if the interaction of replication forks with Tus contributes to the increased mutation rate within this region, we performed an MA experiment on a Δ*tus* <u>Δ</u>*mutL* mutant strain. As shown in Figure 2G, Supplementary Figure S2G, Tables 1 and 2, loss of Tus did not affect the BPS density pattern.

The Ter macrodomain (MD) extends from 1200 Kb to 2200 Kb (12), which is roughly from bin 20 to bin 28 in our figures. The structure of the Ter MD is maintained by the MatP protein, which binds to 23 *matS* sites within this region (41). In the absence of MatP, the Ter MD is disorganized, the DNA is less compact, and the Ter MD segregates too early in the cell cycle and fails to localize properly at midcell (41, 42). Because the mobility of the Ter MD is increased in the absence of MatP, DNA interactions across MD barriers can occur in Δ*matP* mutant cells (41).

As shown in Figure 2H, Supplementary Figure S2H, Tables 1 and 2, loss of MatP caused a severe disruption of the BPS density pattern. The mutation rates in the Ter MD were depressed whereas new peaks appeared on either side of the Ter MD in the Right and Left MDs. Interestingly, the BPS pattern near the origin was maintained in the right but not in the left replichore.

### Recombination and the SOS response

Homologous recombination is intimately connected to replication. As replication proceeds, various blocks, such as DNA lesions, transcription complexes, and DNA secondary structures, can cause the replisome to pause and to eventually disassemble. This potentially lethal event is prevented by recombination, which can repair and restart the replisome (43). In addition, the termination region is subject to hyperrecombination (44, 45) particularly in the region bounded by TerA and TerB (our bins 21 to 24), named the terminal recombination zone (TRZ) (46, 47).

Elimination of *E. coli’s* major recombinase, RecA, had a modest effect on the mutational density pattern. As shown in Figure 3A, Supplementary Figures S3A, S5D, S6D, Tables 1 and 2, in the *recA mutL* and *recA mutS* mutant strains, BPS rates declined in bins 21 to 25, corresponding fairly well to the TRZ described above. However, the hyperrecombination within the TRZ is dependent on the recombination pathway defined by participation of the RecBCD complex. When we eliminated RecB, the mutational density pattern did not phenocopy that seen when RecA was absent (Supplementary Figures S5E, S6E, Tables 1 and 2). Either our protocol is not sensitive enough to detect an effect of loss of RecB, or another recombination pathway, *e.g*. RecFOR (48), is sufficient to maintain the mutation rate in the region.

**Figure 3.**
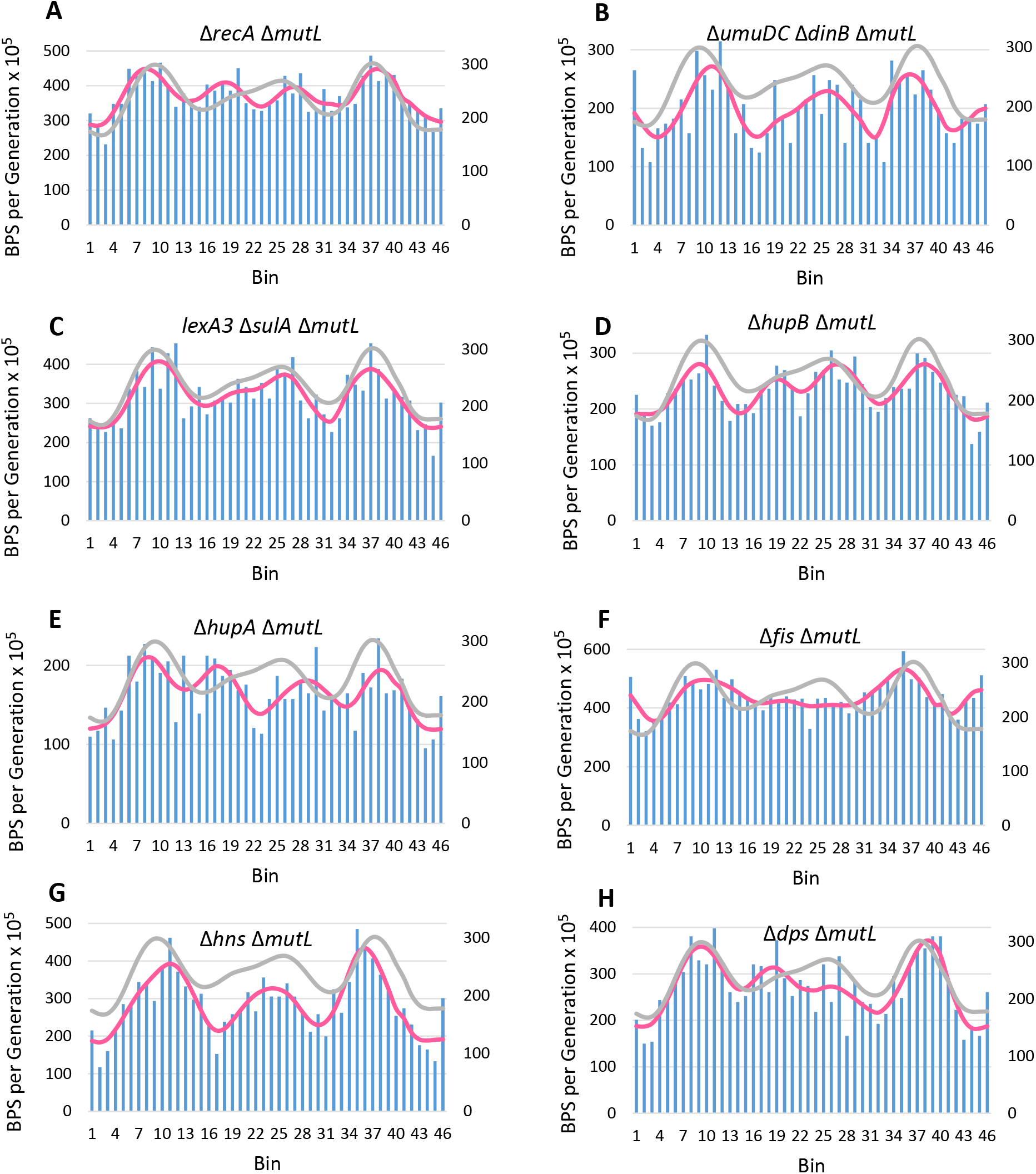
Loss of RecA, HUα, or Fis changes the BPS pattern, but the SOS response and loss of other NAPs do not. For a description of the plots, see the legend to Figure 2. **Figure 3A, B, and C.** Loss of RecA alters the BPS density pattern in the terminus region, but not through its role as a master regulator of the SOS response. RecA (Figure 3A) is *E. coli’s* major recombinase. The *dinB* and *umuDC* genes (Figure 3B) encode the error-prone DNA polymerases, Pol IV and Pol V, that are induced as part of the SOS response. The *lexA3* allele encodes a non-inducible repressor of the SOS genes. Deletion of *sulA* prevents lethal filamentation after induction of the SOS response (not relevant to this study). Strains: 3A, PFM422; 3B, PFM118; 3C, PFM120. **Figure 3D, E, F, G, and H.** Loss of the HUα subunit of HU or of Fis changes the BPS density pattern, but loss of the HUβ subunit of HU, HNS, or DPS has only minor effects. Strains: 3D, PFM259; 3E, PFM258; 3F, PFM317/318; 3G, PFM741/742; 3H, PFM713.

In addition to its role in recombination, RecA is also a master regulator of the SOS response to DNA damage, which includes the induction of two error-prone DNA polymerases, DNA Pol IV and V. To test whether these polymerases are involved in determining the BPS density pattern, we performed an MA experiment on a strain deleted for the genes that encode Pol IV, *dinB*, and PolV, *umuDC*. As shown in Figure 3B,Supplementary Figure S3B, Tables 1 and 2, the BPS density pattern in the *mutL dinB umuDC* mutant strain was not significantly different than the MMR^−^ pattern. The genes of the SOS response are repressed by the LexA protein; the *lexA3* allele encodes a super-repressor LexA protein that prevents the SOS genes from being induced (49). When this allele was present the BPS density pattern was also unaffected (Figure 3C, Supplementary Figure S3C, Tables 1 and 2). Thus, the SOS response appears to play no role in determining the pattern of BPSs across the chromosome.

### Nucleoid Associated Proteins

In a previous report (1), we found that the BPS density pattern of a Δ*mutL* strain was correlated with the density of genes activated by the HU protein and repressed by the Fis protein. Combining these two factors in a linear correlation model accounted for 33% of the variation in the mutational data. HU constrains supercoils and compacts the DNA into nucleosome-like particles; Fis also constrains supercoils but, in addition, bends the DNA (31). While both of these NAPs affect transcription, the general lack of correlation of the BPS rate with transcriptional levels (7); also see above) led us to hypothesize that mutation rates across the chromosome were correlated not with transcription *per se*, but with areas of high DNA structure (1). To further test this hypothesis we preformed MA experiments with MMR^−^ mutant strains also defective for each of a number of NAPs.

HU exists as a dimer of its two subunits, HUα and HUβ, encoded by the paralogous genes *hupA* and *hupB*, respectively, in the three possible configurations. While loss of both subunits confers a severe growth defect, loss of only one has little consequence during a normal growth cycle, suggesting they can substitute for each other. HUαβ is the dominant form over most of the cell cycle, but significant amounts of HUα2 are found during lag phase and early exponential phase, and HUβ2 is prominent in stationary phase (30). Chromatin immunoprecipitation sequencing (ChIP-Seq) results revealed that HU binds non-specifically to the chromosome and the DNA binding patterns of the three dimers appear to be identical (50).

We performed MA experiments with both Δ*hupA* Δ*mutL* and Δ*hupB* Δ*mutL* mutant strains. As shown in Figures 3D, 3E, Supplementary Figures S3D, S3E, Tables 1 and 2, although loss of *hupB* appeared to affect the BPS density pattern, the differences from the MMR^−^ pattern were not significant. However, loss of *hupA* depressed mutation rates across the chromosome, particularly in the terminus region, while creating new peaks on either side of the Ter MD.

Fis has many binding sites across the chromosome, but, based on data at RegulonDB, Sobetzko *et al*. (2012) (51) reported more Fis binding sites in the origin region. However both Chromatin immunoprecipitation plus microarray analysis (ChIP-chip) and ChIP-Seq studies found the density of Fis binding to be more-or-less constant across the chromosome (32, 52). Loss of Fis in both Δ*fis* Δ*mutL* and Δ*fis* Δ*mutS* mutant strains tended to flatten the BPS density pattern across the chromosome except in the region around the origin (Figure 3F, Supplementary Figures S3F, S5F, S6F, Tables 1 and 2).

The NAP HNS binds to DNA at its high-affinity binding sites and then spreads by oligomerization along A:T rich regions of DNA. Bridging between HNS-DNA complexes condenses the DNA into a few clusters per chromosome (53–55). However, as shown in Figure 3G, Supplementary Figure S3G, Tables 1 and 2), loss of HNS had little effect on the BPS density pattern, and, thus, the long-range structures produced by HNS appear not to affect BPS rates.

The DPS protein accumulates in stationary phase cells, condenses the nucleoid into a crystalline-like state, and protects the DNA from oxidative and other damage (56). Despite this radical physical change, loss of DPS had little effect on the BPS density pattern Figure 3H, Supplementary Figure S3H, Tables 1 and 2). Of course, we do not know the degree to which cells in stationary phase contribute to the BPS rates under our experimental conditions.

### Proofreading

As mentioned above, epsilon is the proofreading subunit of the DNA polymerase III holoenzyme. The *mutD5* allele encodes an epsilon protein that is inactive for proofreading, and strains carrying this allele have a mutation rate 4000-fold greater than that of wild-type strains, and 35-fold greater than that of MMR-defective strains (10). As shown in Figure 4A, Supplementary Figure S4A, Tables 1 and 2, when proofreading was inactive but MMR was active, the BPS density pattern was less dramatic than when MMR was inactive and proofreading was active; but, nonetheless, the wave pattern was basically the same. When both MMR and proofreading were inactive, which reveals the mutations solely due to replication errors, the BPS density pattern was nearly flat but around the origin it retained significant correlations to the patterns of both the MMR-defective and *mutD5* mutant strains (Figure 4B, Supplementary Figure S4B, Tables 1 and 2). Thus, the pattern of BPS on both sides of the origin appears to be established by replication errors and then elsewhere across the chromosome the density pattern is largely due to differential error-correction by both proofreading and MMR.

**Figure 4.**
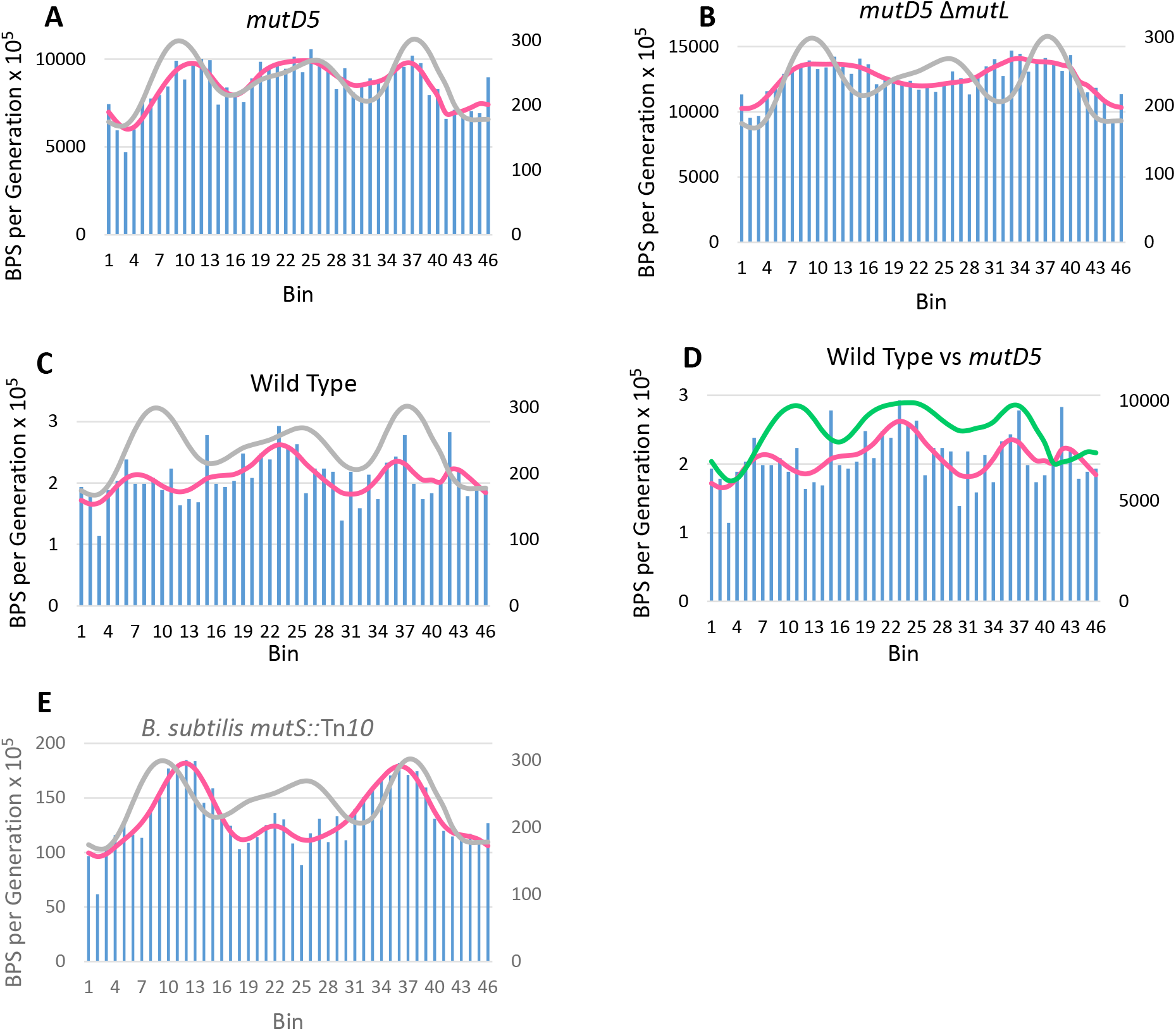
The BPS density pattern is a result of biased correction by replication proofreading and MMR. For a description of the plots, see the legend to Figure 2. **Figure 4A and B.** A strain with deficient proofreading but active MMR yields a BPS pattern similar to a MMR-defective strain, but a strain with neither proofreading nor MMR does not. The *mutD5* allele encodes an exonuclease deficient proofreader. Strains: 4A, PFM163; 4B, PFM165/397/399. **Figure 4C and D.** The BPS density pattern in the wild-type strain does not match the pattern of either the MMR-defective strain, or the *mutD5* mutant strain. The green line in Figure 1D is the Daubechies wavelet transform of the *mutD5* mutant strain (pink line in Figure 4A). Strains: 4C and D, eight strains with wild-type mutational phenotypes (see text). **Figure 4E.** MMR-defective *B. subtilis* also has a symmetrical BPS density pattern, but it is different than *E. coli’s* pattern. Strains: three *B. subtilis* mutant strains with the same mutational phenotype (see supplementary Tables S1 and S3).

### Wild-type

The mutational density wave patterns evident in our data, and in data from other bacteria (2, 3), were obtained when MMR was inactive. Thus, the question arises: does the pattern appear in wild-type strains? It is difficult to answer this question because mutation rates in wild-type strains are so low (in *E. coli*, 120-fold lower than that of MMR-defective strains (7, 11) that enormous experiments would have to be conducted in order to accumulate enough mutations to approach statistical confidence. In a recent study we compared the mutation rates and spectra of a number of *E. coli* strains defective in various DNA repair activities; of these, the results from seven strains were indistinguishable from those of the wild-type parent (57). By combining the mutations from these strains, we achieved 1933 BPSs (11), enough to expect to see a wave pattern if it existed. As is evident in Figures 4C, 4D, Supplementary Figures S4C, S4D, Tables 1 and 2, these BPSs did not create fall into a recognizable pattern. Indeed, the pattern from the wild-type strains appears to be random; the variance to mean ratio of the binned mutations is 1.4, indicating the values are not disperse, and the MatLab “runstest”, a test for runs, returns a P value of 0.58, also indicating that the bin values are random. In the wild-type strain both MMR and proofreading are active, and, while these two activities have similar correction biases (Figure 4A), proofreading is much more powerful, producing the pattern seen in the MMR-defective strains. Although we cannot conclude that the mutational density pattern in the wild-type strain is other than random, it does have similarity to both the patterns seen in the MMR-defective strains and in the *mutD5* mutant strain, particularly around the terminus (Figures 4C, 4D, Supplementary Figures S4C, S4D and Tables 1 and 2).

#### Bacillus subtilis

In additional to *E. coli* strains, symmetrical mutational density patterns have been demonstrated in MMR-derivatives of *Vibrio fischeri, V. cholera* (2–4), *Pseudomonas fluorescens* (5), and *P. aeruginosa* (6). Here we add *Bacillus subtilis* to this list. As shown in Figure 4E, Supplementary Figure S4E, Tables 1 and 2, the BPS mutation rates in a *B. subtilis mutS::Tn10* mutant strain fell into a wave like pattern that was symmetrical about the origin. Although similar in shape, the pattern was significantly different from that of *E. coli* (Figure 4E, Supplementary Figure S4F, Tables 1 and 2). However, as in *E. coli*, the BPS rate appeared to increase in the terminus region, which in *B. subtilis* is not 180° from the origin and corresponds to bins 20-24 in Figure 4E.

## Discussion

In this report we have examined a number of factors that could be responsible for establishment and maintenance of the symmetrical wave-like BPS density pattern across the chromosome. In broad terms these factors were: transcription; DNA replication initiation, progression, and termination; recombination and the SOS response to DNA damage; the binding of nucleoid-associated proteins; and, error-correction by MMR and proofreading. As discussed above, we found that transcription and the SOS response had little effect, and the effect of recombination was modest and confined to the terminal region. We discuss the more significant, factors in greater detail here.

### DNA replication initiation, progression, and termination

Providing additional replication origins, either by eliminating RNase H1 or by inserting an ectopic *oriC* (*oriZ*), did not disrupt the wave (Figure 1C and 1D). However, when *oriZ* was the only origin of replication, the region of depressed BPS rate that surrounds *oriC* when it is in the normal position was re-established about the new origin (Figure 1E). Note that only 5.1 Kb of DNA containing *oriC* was relocated (19), whereas the region of reduced mutation rate is about 200 Kb; thus, the mutation rate is not determined just by the DNA sequence surrounding the origin. We hypothesize that the process of replication initiation protects the DNA from damage and/or newly established replication forks have a low error-rate. In addition, the BPS mutation rate was increased for about 1000 Kb (10 bins) on either side the new origin so that it resembled the same region about the normal origin. The size of this area is close to the same size detected as “interacting zones” around *oriZ* (58). However, the mutational density pattern across the rest of the chromosome did not re-establish itself to be symmetrical about *oriZ*, but remained symmetrical about the absent *oriC*. Thus, other factors must be important at distant regions. We can also conclude that the overall structure of the BPS pattern across the chromosome is not determined by active replication initiation *per se*, but may have evolved in response to replication initiation.

Both *V. fischeri* and *V. cholera* have multiple circular chromosomes of different sizes; by comparing the mutational density patterns of these chromosomes, Dillon et al, 2018 (2) identified the timing of replication as a significant determinant of the mutational density patterns. As mentioned above, they suggested that the pattern could be the result of variations in the levels of dNTPs as origins fire during rapid growth. Our results provide partial support for this hypothesis, but also indicate that growth on rich medium, not growth rate *per se*, is a significant determinant of the mutational density pattern, possibly because of effects on the expression and DNA binding of HU and Fis (see below). Our results with a Δ*nrdR* mutant strain (Figure 2E) also show that dNTP levels are important, but, again, other factors are driving the overall wave pattern. In addition, the results with the Δ*rep* mutant strain (Figure 2F) indicate that radically interrupting the progression of the replication fork disrupts the mutational density pattern.

After declining to a local minimum about 3/5^th^ of the distance along each replichore, mutation rates rise in the terminus region (Figure 1A). We originally hypothesized that this increase was due to collisions of the replication complexes with the Tus anti-helicase, most of which would take place in the region between bins 18 and 31 (1). However, elimination of Tus had no effect on the wave pattern (Figure 2G, Supplementary Figure S2G), refuting this hypothesis. But, eliminating MatP, the protein that maintains the structure of the Ter MD, reduced mutation rates in the Ter MD, suggesting that replication of highly structured DNA is error-prone (Figure 2H, Supplementary Figure S2H). Interestingly, in Δ*matP* mutant cells mutation rates increased in the Right and Left MCs, suggesting that in the absence of MatP, these adjacent regions of DNA gained structure. The mechanism by which MatP structures the Ter MC is not clear (59, 60), but it probably involves supercoils which, when unconstrained, could migrate into adjacent areas. However, two studies have found that loss of MatP increases the mobility and long-range interactions of adjacent DNA (41, 61). Resolution of these conflicts will require further experimentation.

#### Nucleoid-associated proteins

Our previous results predicted that the NAPs HU and Fis should play a role in establishing or maintaining the wave-pattern of BPS, but HNS should not (1). The results presented here confirmed that prediction. In addition, we also found that Dps had no effect on the BPS density pattern. Because the NAPs affect gene expression in various ways, we cannot conclude that DNA binding by the NAPs themselves is responsible for the mutational pattern. And, indeed, we found no significant positive correlations between the BPS pattern and published binding sites of the NAPs, although the location of the binding sites themselves vary widely among published results (e.g. see (32, 50) and the data in RegulonDB (62). Nonetheless, we favor the hypothesis that structuring of the DNA by the NAPs, directly or indirectly, contributes to the BPS pattern.

The local effect of HU binding is to bend the DNA, but dimer-dimer interactions produce higher-order HU-DNA complexes that can constrain negative supercoils (63, 64). The analysis of the effect of HU on the mutational density pattern is also complicated by the variation in the cellular concentrations of the three forms with growth cycle (30). Although the two subunits can compensated for each other for viability, the affinities of three forms of HU for various DNA structures (linear, nicked, and gapped) differ (65). Both HUα_2_ and HUαβ can constrain supercoils, but HUβ2 apparently cannot, at least *in vitro* (30). Since HUβ is not stable (66), the phenotype of a *ΔhupA* mutant strain may simply be that of low cellular HU concentration.

While loss of HUβ had little effect on the mutational density pattern (Figure 3D, Supplementary Figure S3D), loss of HUα reduced the overall BPS rate by 33% and also changed the pattern (Figure 3E, A Supplementary Figure S3E). In the Δ*hupA* Δ*mutL* mutant strain, mutation rates were low in the terminus region but elevated in the Right and Left MCs, very similar to the pattern seen in the Δ*matP* Δ*mutL* mutant strain (Figure 2H). The low mutation rate in the Ter MD corresponds well with the density of genes activated in wild-type cells but not in HU-deleted cells (67). HU regulates transcription by modifying the DNA superhelical density, suggesting that, like MatP, HU increases the structure of the DNA in the terminal region, causing DNA replication to become error-prone.

The mutational pattern in the Δ*hupA* Δ*mutL* mutant strain is also similar to that of the Δ*recA* Δ*mutL* strain (Figure 3A). An mutation accumulation experiment with a Δ*hupA* Δ*recA* Δ*mutL* mutant strain resulted in a pattern similar to both single mutant strains, and so was not informative (data not shown).

Fis is a major transcriptional regulator, either activating or repressing, directly or indirectly, nearly a thousand genes (52). At some promotors Fis acts as a classic transcriptional regulator by interacting with RNA polymerase, but at other promotors Fis affects gene expression by altering the DNA superhelical density (68). The Fis DNA binding sequence is degenerate and the estimated number of binding sites identified by sequence analysis across the genome varies widely among studies. The results of ChIP analyses also vary: the number of DNA regions associated with Fis in different studies ranges from 200 to 1500, which may reflect the effects of different growth conditions (32, 52, 69).

The loss of Fis nearly doubled the overall BPS rate, but flattened the wave pattern outside of the origin region (Figure 3F, Supplementary Figures S5F, S6F). As mentioned above, the mutational density pattern MMR^−^ strains is correlated to the density of genes activated in a Δ*fis* mutant strain as reported by Blot *et al*, 2006 (70). This correlation is particularly strong in bins 14 to 33 (ρ = 0.62, P = 0.003), which corresponds to the area flattened in the Δ*fis* Δ*mutL* and Δ*fis* Δ*mutS* mutant strains. However, no correlation exists between our mutational data and genes found to be responsive to Fis in a recent study (32). Clearly more studies are needed to resolve these conflicts.

#### Error-correction

Assuming that the BPS recovered from the *mutD5* Δ*mutL* mutant strain are due to intrinsic errors made by DNA polymerase, we conclude that the polymerase is accurate close to the origin, then becomes increasingly less accurate as replication proceeds to about 1/3 of the replichore, at which point accuracy increases again (Figure 4B). The mutational pattern in the MMR-defective strains must reflect the biased ability of proofreader to correct these polymerase errors. Thus, proofreading is effective close to the origin, but then declines for about 1/3 of the replichore, increases in the Right and Left MCs, but declines again in the terminus region (Figures 1A, 4B). Mutations in the *mutD5* mutant strain are due to the failure of MMR to correct the polymerase errors that survive in the absence of proofreading. Since the mutational pattern in the *mutD5* mutant strain mimics that of the MMR-defective strains (Figure 4A, Supplementary Figure S4A, Tables 1 and 2), error correction by MMR, although much less powerful, apparently has the same biases as error correction by proofreader.

Given the above considerations, we would expect the mutational density pattern in the wild-type parental strain to mimic that of the MMR-defective strains. However, the nearly 2000 BPS accumulated in eight strains with wild-type mutational phenotypes appear randomly distributed across the chromosome (Figures 4C, 4D). One explanation for this discrepancy is that when both proofreading and MMR are active, the mutation rate is reduced to the point that other, weaker DNA repair and mutagenic activities obscure the underlying pattern. Although not significantly correlated, the wild-type BPS pattern is similar to the MMR^−^ pattern, particularly around the terminus, supporting this hypothesis. Alternatively, although the eight strains that we combined to produce 2000 BPS have the same mutation rates, they may have different wave patterns that tend to negate each other and obscure the underlying wild-type pattern.

## Conclusions

By applying mutation accumulation followed by whole-genome sequencing to bacterial strains mutant in various activities we have determined that the most important factors determining the symmetrical pattern of BPS rates across the chromosome are: the initiation, progression, and termination of DNA replication, replication error-correction, and chromosome structure. Because of the conservation of these factors, our results should apply to most bacteria, and possibly eukaryotes, and imply that different regions of the genome evolve at different rates.

## Methods

### Bacterial Strains and Media

The strains used in this study are listed in Supplementary Table S1 and the methods of their construction are in Supplementary file: Materials and Methods. Standard media and antibiotics were used (see Supplementary file: Material and Methods).

### Mutation accumulation experiments

The complete MA procedure has been extensively described (7, 11, 57). Details are given in the Supplementary file: Material and Methods

### Genomic DNA Preparation, Library construction, Sequencing, and Sequence Analysis

Details are given in the Supplementary file: Material and Methods.

### RNA sequencing

The *E. coli* strain PFM144, which is PFM2 Δ*mutL* (11), was grown in LB and aliquots collected during lag (OD = 0.022), log (OD = 0.3), and stationary (OD = 1.5) phase. The number of cells collected was kept constant for each growth phase. Total RNA was extracted, DNA and rRNA were removed, and libraries constructed and sequenced as detailed in the Supplementary file: Material and Methods. Three biological triplicates were prepared for each growth phase.

A complete analysis of the RNA-Seq results will be the subject of a subsequent report. For this report the numbers of RNA-Seq reads for each condition were first normalized to the number of reads mapped to the gene *holD*, which was determined by rtPCR to be expressed at the same level in all phases of growth. The means of the normalized RNA reads from the triplicates were then binned into the same bins used for the mutational analysis. A fourth-order Daubechies wavelet transform was performed on the binned RNA-Seq reads as described for the mutational data (1).

### Statistical Analysis

To obtain the BPS density patterns, the numbers of BPSs were binned into 46 bins, each 100 Kb long, as described (1). A fourth-order Daubechies wavelet transform was performed on the binned mutation data as described (1). For presentation in the figures, these results were converted into rates by dividing the number of BPS by the appropriate number of generations. Pearson’s product-moment correlation coefficient, ρ, was used to evaluate the correlations between the binned BPS data (Table 1). Spearman’s nonparametric correlation coefficient was also computed for a few data sets, but gave similar results. To account for multiple comparisons, p values were adjusted using the Benjamini-Hochberg method (71) with the false discovery rate set at 25%, implemented with the MatLab R2018a “mafdr” command. Because comparisons using the data from the same strain are not independent, this adjustment was made separately for each column in Tables 1 and 2.

To further compare the BPS density patterns between two data sets, wavelet coherence was calculated and plotted using the MatLab R2018a “wcoherence” command. While the Daubechies wavelet provides a good visual representation of the binned data, it is not continuous and thus not easily adapted for wavelet coherence analysis. The MatLab program first converts the binned data to Morlet wavelets and then computes the coherence between two of these wavelets. We chose to analyze the data with wavelet coherence because it gives a measure of the correlation between the signals (displayed as colors in the figures) (72, 73). In addition, the MatLab wavelet coherence plot indicates, as a dashed curve, the ‘cone of influence’ within which results are free of artifactual edge effects (72). The relative phase-lag between the two signals is indicated by small arrows: arrows pointing right indicate in-phase, arrows pointing left indicate 180° out-of-phase, and arrows pointing in other directions indicate the various degrees in between. Because the MatLab program assumes a frequency-time series, the X axis of the plot is cycles/sample and the Y axis is time; we converted these to bins/cycle (on an inverted scale) and bins, respectively.

## Declarations

### Ethics approval and consent to participate

Not applicable

### Consent for publication

Not applicable

### Availability of data and material

The sequences and SNPs reported in this paper have been deposited with the National Center for Biotechnology Information Sequence Read Archive https://trace.ncbi.nlm.nih.gov/Traces/sra/ (accession no. in progress) and in the IUScholarWorks Repository (hdl.handle.net/2022/20340). Bacterial strains are available upon request

### Competing interests

The authors declare no competing interests

### Funding

This research was supported by US Army Research Office Multidisciplinary University Research Initiative (MURI) Award W911NF-09-1-0444 to P.L.F. and H.T., the National Institutes of Health T32 GM007757 to B.A.N., and the US Army Undergraduate Research Apprenticeship Program to J. H. and S. R

### Authors’ contributions

B.A.N. and P.L.F. designed the research, analyzed the data, and wrote the paper. B.A.N executed the experiments. W.M., H.L., and H.T. performed the bioinformatic analyses. P.L.F. and H.T. supplied resources. All authors read and approved the final manuscript.

## Acknowledgements

We thank H. Bedwell-Ivers, C. Coplen, M. Durham, J. Eagan, N. Gruenhagen, J. A. Healy, N. Ivers, C. Klineman., H. Lee, E. Popodi, I. Rameses, S. Riffert, H. Rivera, D. Simon, K. Smith, J. Townes, L. Tran, and L. Whitson for technical help. Bacterial strains were kindly provided by R. Schaaper, R. Reyes-Lamothe, D. Kearns, M. Konkol, and The National BioResource Project at the (Japanese) National Institute of Genetics. We also thank S. E.Bell and X. Wang for technical help and discussions.

## Supplemental Material and Methods

### Bacterial strains and media

The strains used in this study are listed in Supplemental Table S1. All E. coli strains were derived from PFM2 (1) or AB1157 (2). The *oriC^+^ oriZ^+^* and Δ*oriC oriZ^+^* strains were a gift from Rodrigo Reyes-Lamothe (McGill University). The *mutD5* allele was obtained from Roel Schaaper (NIEHS). The deletion mutations originated in the Keio collection (3) and were moved by P1 phage transduction (4); the Kn^r^ element was removed by using FLP recombination (5). The deletions were confirmed by PCR analysis using the oligonucleotides listed in Supplemental Table S2. The *B. subtilis* strains were derived from the undomesticated ancestral strain NCIB3610 and were a gift from M.A. Konkol and D.B. Kearns (Indiana University).

Rich medium was Miller Luria Broth (LB) (Difco; BD); minimal medium was Vogel-Bonner minimal medium (VB min) with 0.2% glucose (6). When required, antibiotic concentrations were: carbenicillin (Carb), 100 μg/ml; kanamycin (Kn), 50 μg/ml; nalidixic acid (Nal), 40 μg/ml; chloramphenicol (Cam), 30 μg/ml; and, rifampicin (Rif), 100 μg/ml. Half of these concentrations were used in minimal medium.

### Estimation of mutation rates from fluctuation assays

Mutation rates were determined as described (7), using mutation to Nal^R^ or Rif^R^. The Ma-Sandri-Sarkar maximum likelihood method was used to calculate the mutation rates by using the FALCOR web tool found at www.mitochondria.org/protocols/FALCOR.html (8).

### Mutation accumulation experiments

The MA procedure has been described (1, 9, 10). The MA lines originated from single colonies isolated from a founder colony, obtained by streaking from a freezer stock onto agar plates of the medium to be used in the MA experiment. After incubation overnight at the experimental temperature, one well isolated colony was excised from the agar plate, soaked for 30 minutes in 0.85% NACL + 0.01% gelatin, and then vortexed for 60 seconds. Appropriate dilutions for obtaining well-isolated colonies were then plated onto the appropriate agar plates at the appropriate temperature to start MA lines. Plates were incubated at 37°C for most experiments, or at 28°C for the experiment at low temperature. Each MA line was periodically streaked for a single colony: on LB and supplemented VB min agar plates at 37°C, this was done daily; on VB min and diluted LB agar plates at 37°C, and on LB plates at 28°C, this was done every 48 hours. The number of passes required was determined by the preliminary mutation rate obtained from a fluctuation assay. The parameters of the MA experiment including the number of lines used for each strain and the total number of generations per experiment are given in Supplemental Table S3.

### Estimation of generations

The method to estimate that number of generations undergone in each MA experiment is described (1, 9). The diameter of the single colonies streaked was recorded daily, and then the number of cells in colonies of different diameters was determined for each experiment as described (1). The daily colony diameters were converted to generations and the results summed.

### Genomic DNA preparation, library construction, and sequencing

Genomic DNA was isolated using PureLink Genomic DNA purification kit (Invitrogen) from a predetermined amount of overnight LB culture inoculated from the freezer stocks made after the last passage of the MA line. DNA concentration was measured on an Epoch Microplate Spectrophotometer (BioTek Instruments, Inc.). The identity of the lines was confirmed before library construction by verifying the presence of the expected gene deletions in the gDNA with diagnostic PCR using the oligonucleotides in Supplemental Table S2. Libraries were made by the Indiana University Center for Genomics and Bioinformatics and were sequenced using the Illumina HiSeq 2500 platform at the University of New Hampshire Hubbart Center for Genome Studies or the Illumina NextSeq platform at the Indiana University Center for Genomics and Bioinformatics.

### Sequence analysis

SNP calling is described (1). The reference genome used for *E. coli* was NCBI reference sequence NC_000913.2, and for *B. subtilis* was GenBank: CP020102.1. The Illumina reads were aligned to the referenced genomes using the Burrows-Wheeler short-read alignment tool, BWA version 0.7.9 (11).

Poor sequence coverage resulted in some MA lines being eliminated. Cross-contamination can occur during streaking, resulting in lines with identical mutations. If two lines shared over 50% of their mutations then one of the lines was dropped from further analysis. If lines shared less than 50%, then each shared mutation was assigned to one of the lines and dropped from the others. If lineage could be established the shared mutation was assigned accordingly, otherwise the mutation was assigned randomly.

### RNA sequencing

The *E. coli* strain PFM144, which is PFM2 Δ*mutL* (10), was grown in LB and aliquots collected during lag (OD = 0.022), log (OD = 0.3), and stationary (OD = 1.5) phase. The number of cells collected was kept constant for each growth phase. Cells were pelleted at 10,000gs for 10 min, 1ml of medium was added and the cells were pelleted again at 10,000gs for 2 min. The pellets were then flash frozen in liquid nitrogen and stored at −80°. Three biological triplicates were prepared for each growth phase.

RNA was extracted using FastRNA Pro Blue kit (MP Biomedicals). DNA was removed by using TurboDNase (Ambion). DNA removal was confirmed with diagnostic PCR using primer pair “fis forward” and “fis reverse” (Supplemental Table S2). rRNA was removed using RiboMinus magnetic beads kit (Invitrogen). RNA concentration and purity was assessed with an Epoch Microplate Spectrophotometer (BioTek Instruments, Inc.). Libraries were made by the Indiana University CGB and were sequenced at UT Health Science Center on Illumina HiSeq 2500 platform.

**Table S1.** Bacterial strains used in this study

| Strain | Relevant genotype | Donor | Recipient | Target gene | Reference |
| --- | --- | --- | --- | --- | --- |
| PFM2 | MG1655 *rph*<sup>+</sup> |  |  |  | (1) |
| PFM5 | Δ*mutL* | | | | (1) |
| WX320 | Δ*oriC* *oriZ*<sup>+</sup> (AB1157) | | | | (12) |
| WX340 | *oriC*<sup>+</sup> *oriZ*<sup>+</sup> (AB1157) |  |  |  | (12) |
| PF11 | AB1157 |  |  |  | Lab strain |
| PFM118 | Δ*umuDC* Δ*dinB* Δ*mutL* | | | | (10) |
| PFM120 | *lexA3* Δ*sulA* Δ*mutL* | | | | This Study |
| PFM163 | *mutD5* |  |  |  | (13) |
| PFM165/397/399 | *mutD5* Δ*mutL* | | | | (13) |
| PFM244 | Δ*mutL*, *scarless* | | | | (10) |
| PFM256 | Δ*tus* Δ*mutL* | JW1602 | PFM244 | Δ*tus*::Kn<sup>R</sup> | This Study |
| PFM257 | Δ*matP* Δ*mutL* | JW0939 | PFM244 | Δ*matP*::Kn<sup>R</sup> | This Study |
| PFM258 | Δ*hupA* Δ*mutL* | JW3964 | PFM244 | Δ*hupA*::Kn<sup>R</sup> | This Study |
| PFM259 | Δ*hupB* Δ*mutL* | JW0430 | PFM244 | Δ*hupB*::Kn<sup>R</sup> | This Study |
| PFM317/ 318 | Δ*fis* Δ*mutL* | JW3229 | PFM244 | Δ*fis*::Kn<sup>R</sup> | This Study |
| PFM342/343 | Δ*mutS*, *scarless* | | | | (10) |
| PFM421 | Δ*rnhA* Δ*mutL* | JW0204 | PFM244 | Δ*rnhA*::Kn<sup>R</sup> | This Study |
| PFM422 | Δ*recA* Δ*mutL* | JW2669 | PFM244 | Δ*recA*::Kn<sup>R</sup> | This Study |
| PFM424 | *ΔrecA ΔmutS* | JW2669 | PFM342 | *ΔrecA::Kn<sup>R</sup>* | This Study |
| PFM426 | *ΔoriC oriZ<sup>+</sup> ΔmutL*<br>(AB1157) | JW4128 | WX320 | *ΔmutL::Kn<sup>R</sup>* | This Study |
| PFM430/ 431 | *oriC<sup>+</sup> oriZ<sup>+</sup> ΔmutL*<br>(AB1157) | JW4128 | WX340 | *ΔmutL::Kn<sup>R</sup>* | This Study |
| PFM456 | *ΔrecB ΔmutL* | JW2788 | PFM244 | *ΔrecB::Kn<sup>R</sup>* | This Study |
| PFM482 | *Δfis ΔmutS* | JW3229 | PFM343 | *Δfis::Kn<sup>R</sup>* | This Study |
| PFM533/ 534 | *ΔseqA ΔmutL* | JW0674 | PFM244 | *ΔseqA::Kn<sup>R</sup>* | This Study |
| PFM661 | *ΔhupA ΔrecA ΔmutL* | JW2669 | PFM258 | *ΔrecA::Kn<sup>R</sup>* | This Study |
| PFM669 | *ΔmutL* (AB1157) | JW4128 | PF11 | *ΔmutL::Kn<sup>R</sup>* | This Study |
| PFM677 | *Δrep ΔmutL* | JW5604 | PFM244 | *Δrep::Kn<sup>R</sup>* | This Study |
| PFM713 | *Δdps ΔmutL* | JW0797 | PFM244 | *Δdps::Kn<sup>R</sup>* | This Study |
| PFM741 | *Δhns ΔmutL* | JW1225-2 | PFM244 | *Δhns::Kn<sup>R</sup>* | This Study |
| PFM799 | *ΔnrdR ΔmutL* | JW0403 | PFM244 | *ΔnrdR::Kn<sup>R</sup>* | This Study |
| DK2140/2141/<br>2142/2143 | *Bacillus subtilis mutS::Tn10* |  |  |  | M. Konkol & D.<br>Kearns, personal<br>communication |

In all strains the Kn<sup>R</sup> element was removed by FLP recombination (5).
Scarless: Scarless gene deletion using a cat-I-SceI cassette (14).

**Table S2.**
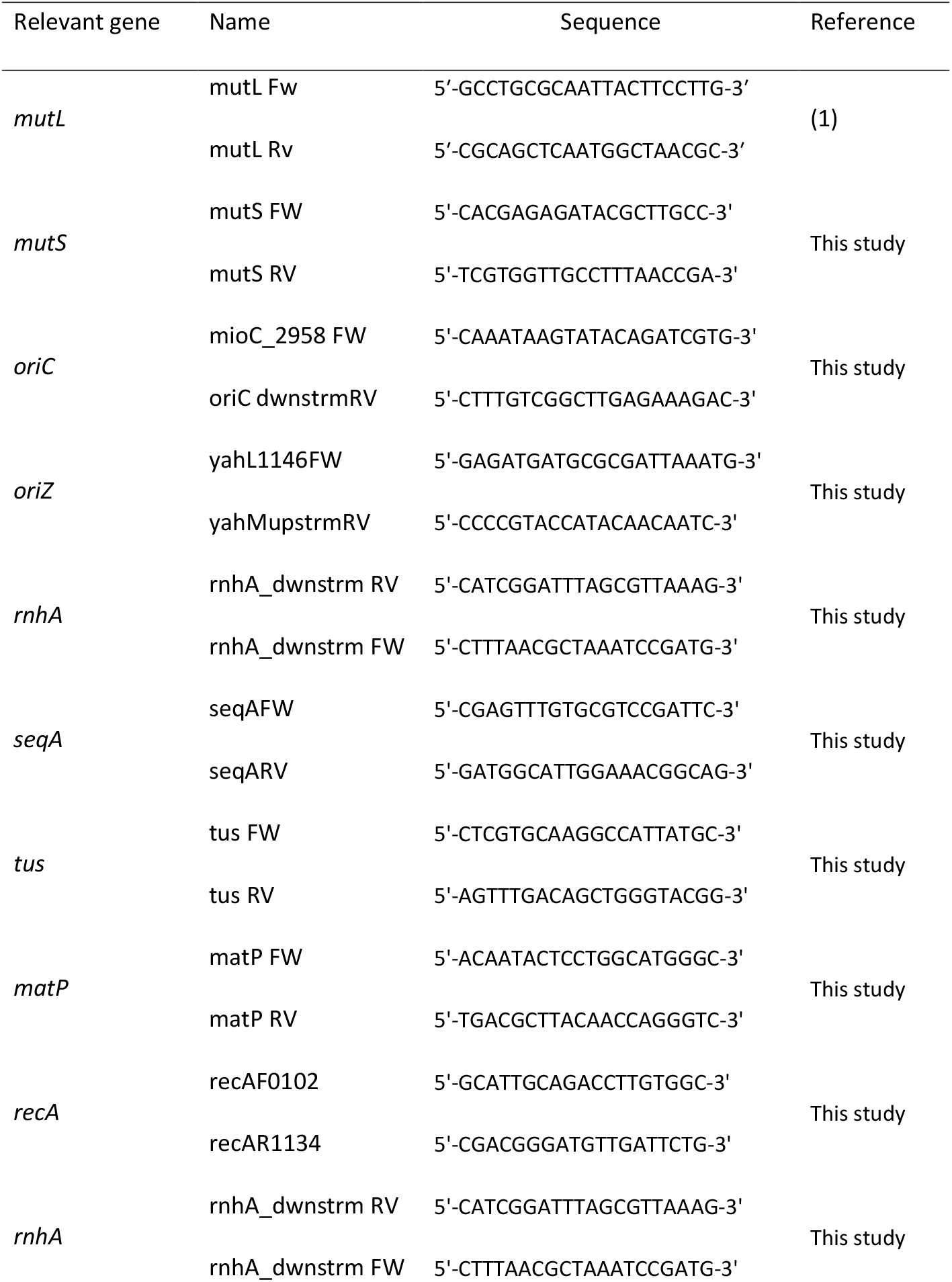
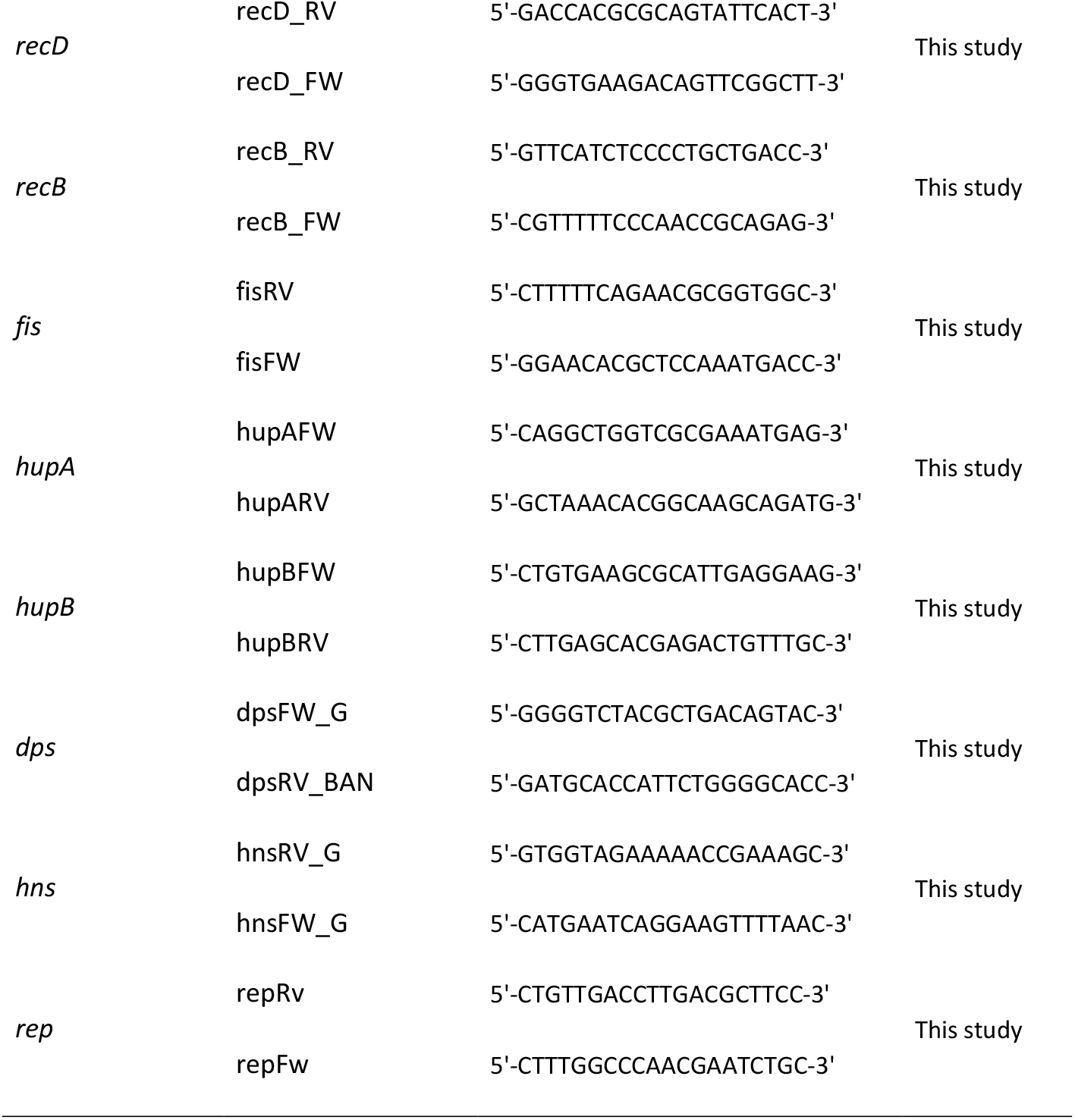
Oligonucleotides used in this study

**Table S3.**
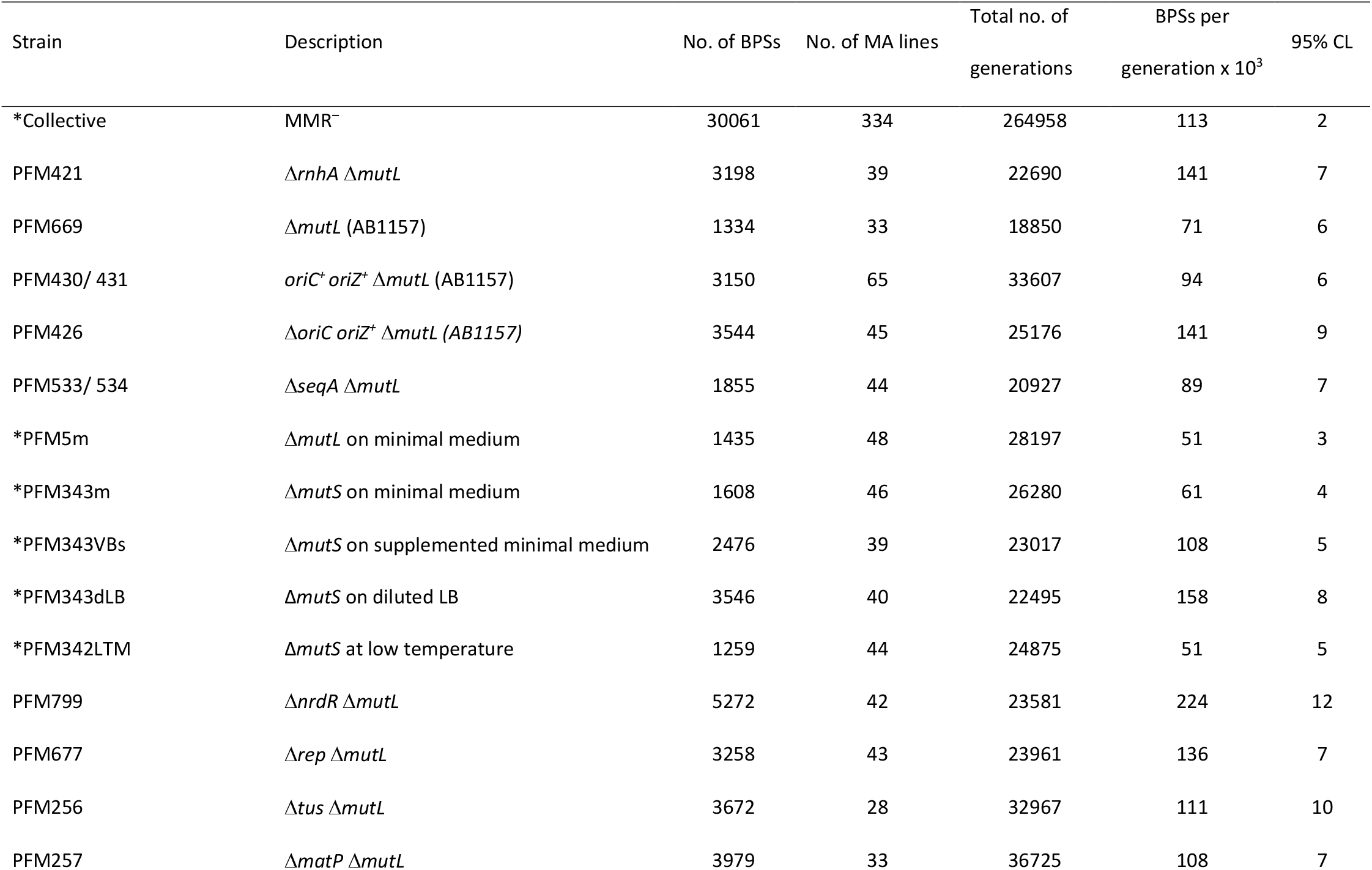
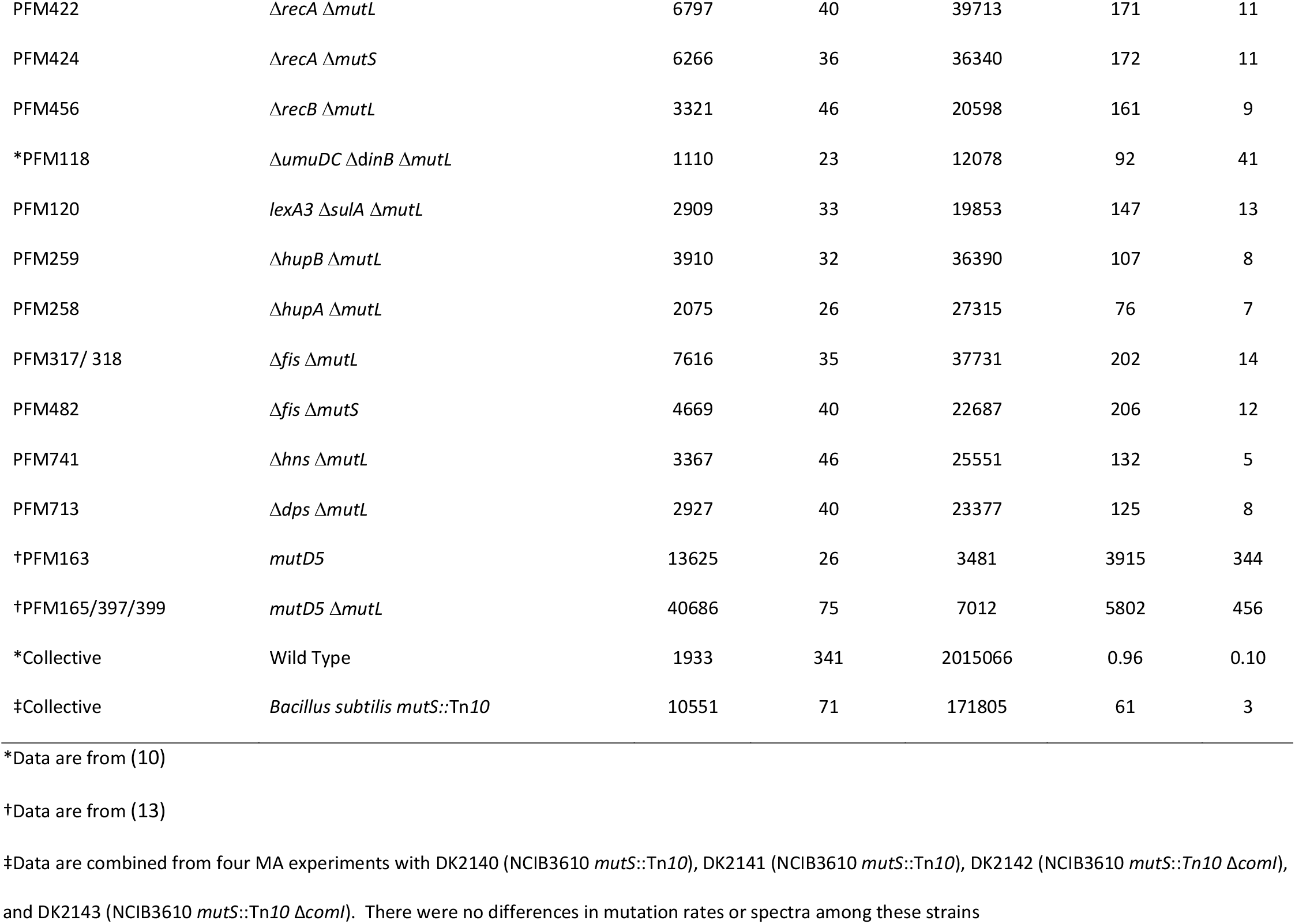
Experimental data

## Supplementary Figure Legends

**Supplementary Figure S1.**
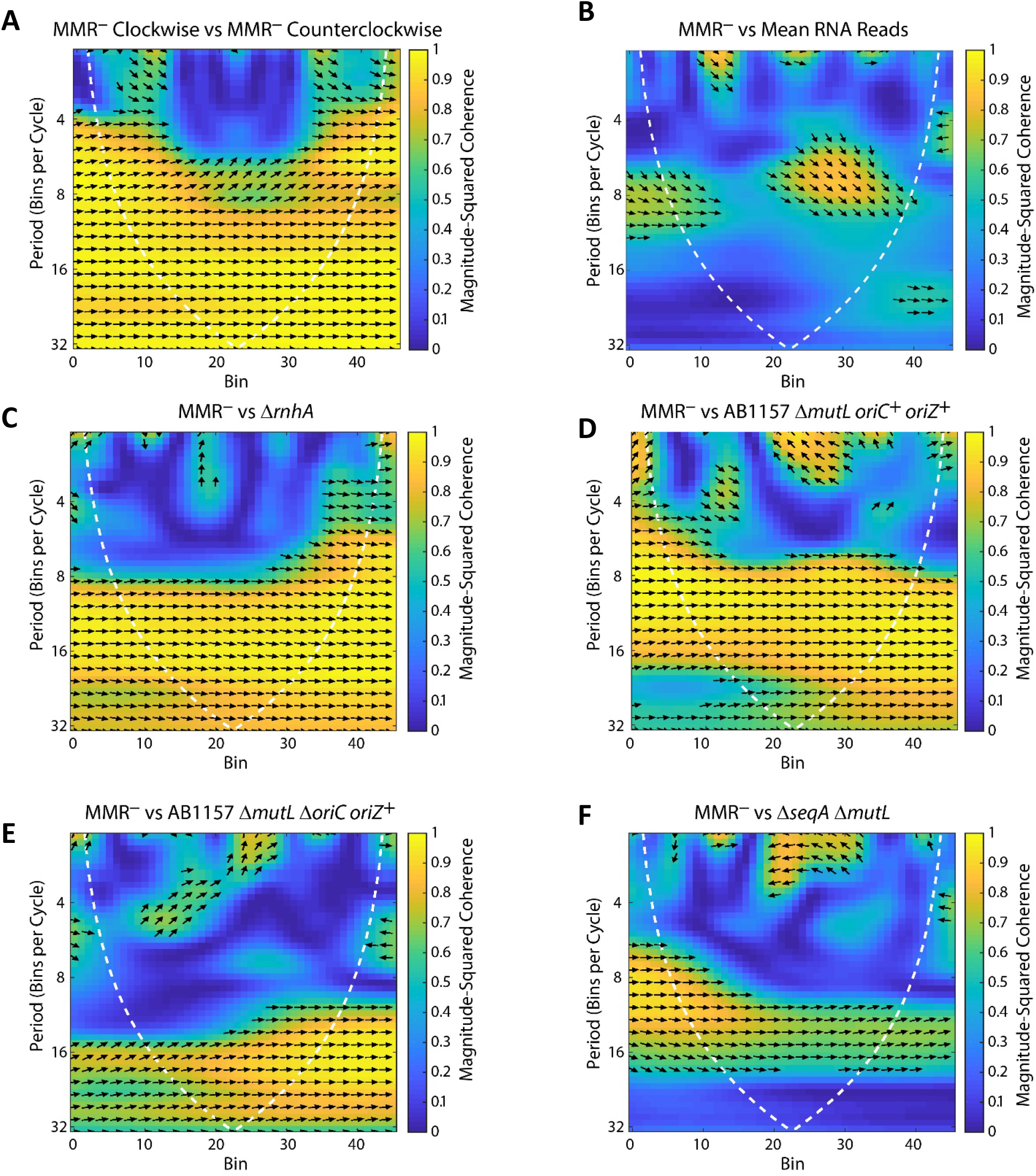
The BPS density pattern is not due to transcription but is affected by moving the origin of replication. Shown are plots of the wavelet coherence between two binned BPS data sets. The colors give the magnitude-squared coherence, a measure of the correlation between the data sets, according to the scale on the left. The dotted line gives the “cone of influence” within which the results are free of artifactual edge effects. Arrows indicate the phase-lag between the two data sets; arrows pointing right indicate in-phase, arrows pointing left indicate 180° out-of-phase, and arrows pointing in other directions indicate the various degrees in between. **Supplementary Figure S1A.** The wavelet coherence between the binned BPS data from the MMR-defective strains taken in the clockwise and counterclockwise directions around the chromosome. This presentation illustrates the almost complete symmetry of the BPS density pattern. **Supplementary Figure S1B.** Plot of the wavelet coherence between the BPS data from the MMR-defective strains and the mean binned reads from RNA-Seq samples taken during lag, log, and stationary growth phases of the Δ*mutL* mutant strain, PFM144. **Supplementary Figures S1C, D, E, and F.** Plots of the wavelet coherence between the binned BPS data from the MMR-defective strains and the indicated mutant strains. See text for a description of the strains. Strains: S1C, PFM421; S1D, PFM430/431; S1E, PFM426; S1F, PFM533/534.

**Supplementary Figure S2.**
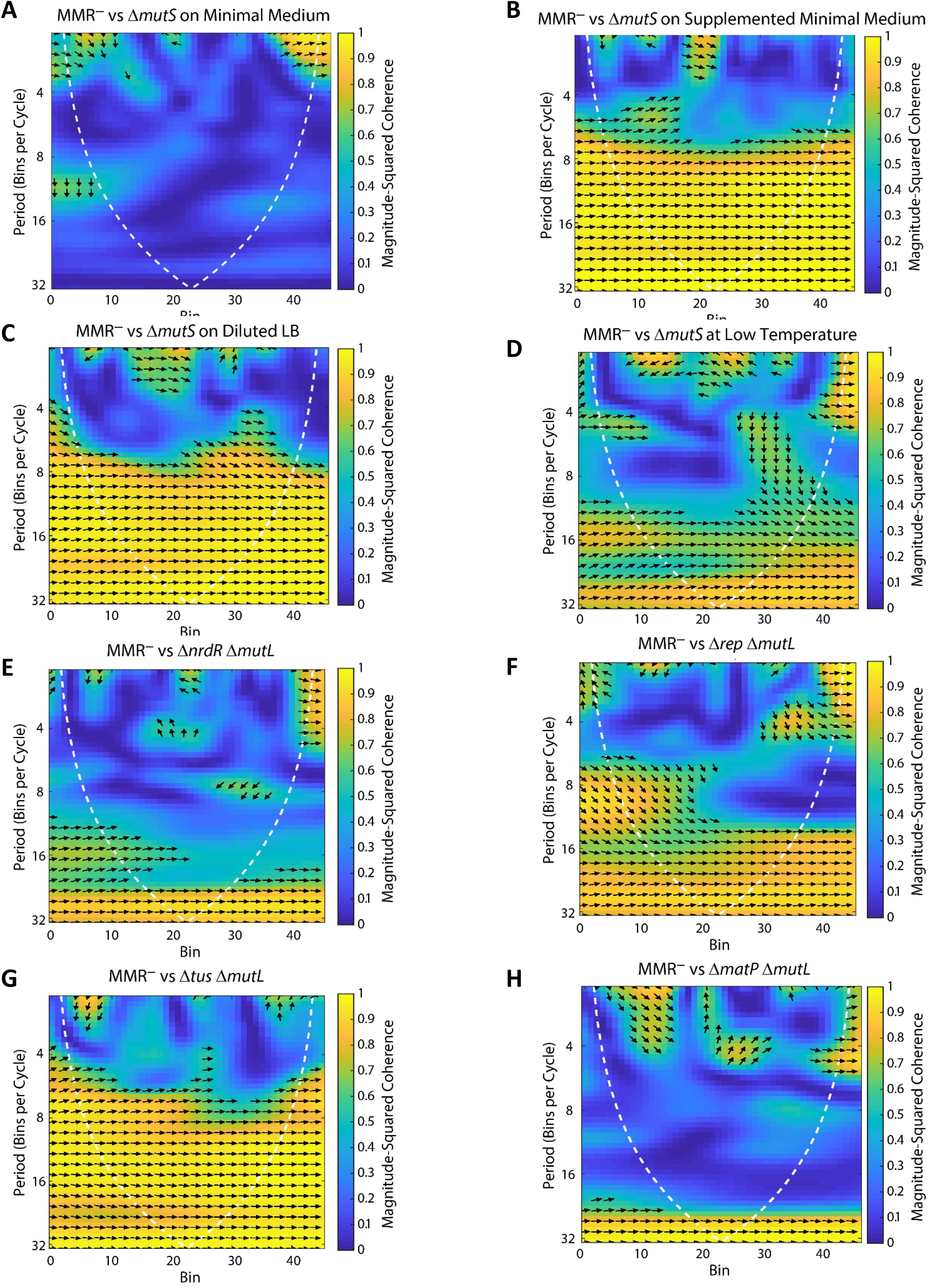
Altering the progression of DNA replication affects the BPS density pattern. Plots of the wavelet coherence between the binned BPS data from the MMR-defective strains and the indicated mutant strains. The colors give the magnitude-squared coherence, a measure of the correlation between the data sets, according to the scale on the left. The dottedline gives the “cone of influence” within which the results are free of artifactual edge effects. Arrows indicate the phase-lag between the two data sets; arrows pointing right indicate inphase, arrows pointing left indicate 180° out-of-phase, and arrows pointing in other directions indicate the various degrees in between. **Supplementary Figures S2A, B, C, and D.** Altering the cellular growth rate by growing a Δ*mutS* mutant strain on different medium and at different temperatures differentially affects the BPS density pattern. In the experiments shown in Plot A, C, and D, the cells grew at about half the normal rate, whereas in the experiment shown in plot B they grew at the normal rate. Strains: S2A, B, and C, PFM343; S2D, PFM342. **Supplementary Figures S2E and F.** Dysregulation of dNTP levels and loss of Rep, the auxiliary replication helicase, affect the BPS density pattern. Strains: S2E, PFM799; S2F, PFM677. **Supplementary Figures S2G and H.** Loss of the Tus antihelicase has no effect, but loss of the terminus organizing protein, MatP, changes the BPS density pattern across the chromosome. Strains: S2G, PFM256; S2H, PFM257.

**Supplementary Figure S3.**
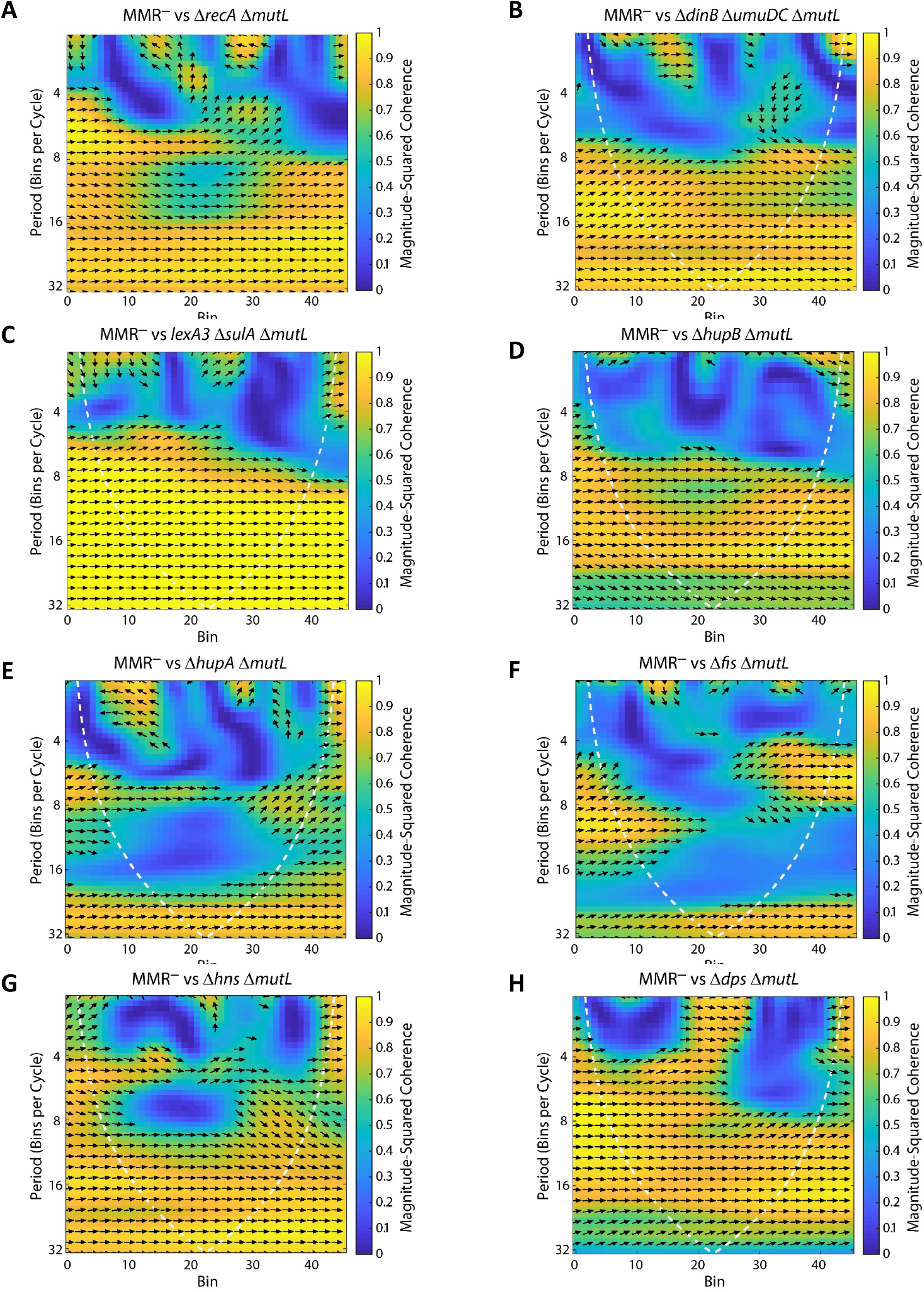
Loss of RecA, HUα, or Fis changes the BPS pattern, but the SOS response and other NAPs do not. For a description of the plots, see the legend to Supplementary Figure S2. **Supplementary Figures S3A, B, and C.** Loss of RecA alters the BPS density pattern in the terminus region, but not through its role as a master regulator of the SOS response. RecA (Figure S3A) is *E. coli’s* major recombinase. The *umuDC* and *dinB* genes (Figure S3B) encode error-prone DNA polymerases that are induced as part of the SOS response. The *lexA3* allele encodes a non-inducible repressor of the SOS genes. Deletion of *sulA* prevents lethal filamentation, which is not relevant to this study. Strains: S3A, PFM422; S3B, PFM118; S3C, PFM120. **Supplementary Figures S3D, E, F, G, and H.** Loss of the HUα subunit of HU or of Fis changes the BPS density pattern, but loss of the HUβ subunit of HU, HNS, or DPS have only minor effects. Strains: S3D, PFM259; S3E, PFM258; S3F, PFM317/318; S3G, PFM741/742; S3H, PFM713.

**Supplementary Figure S4.**
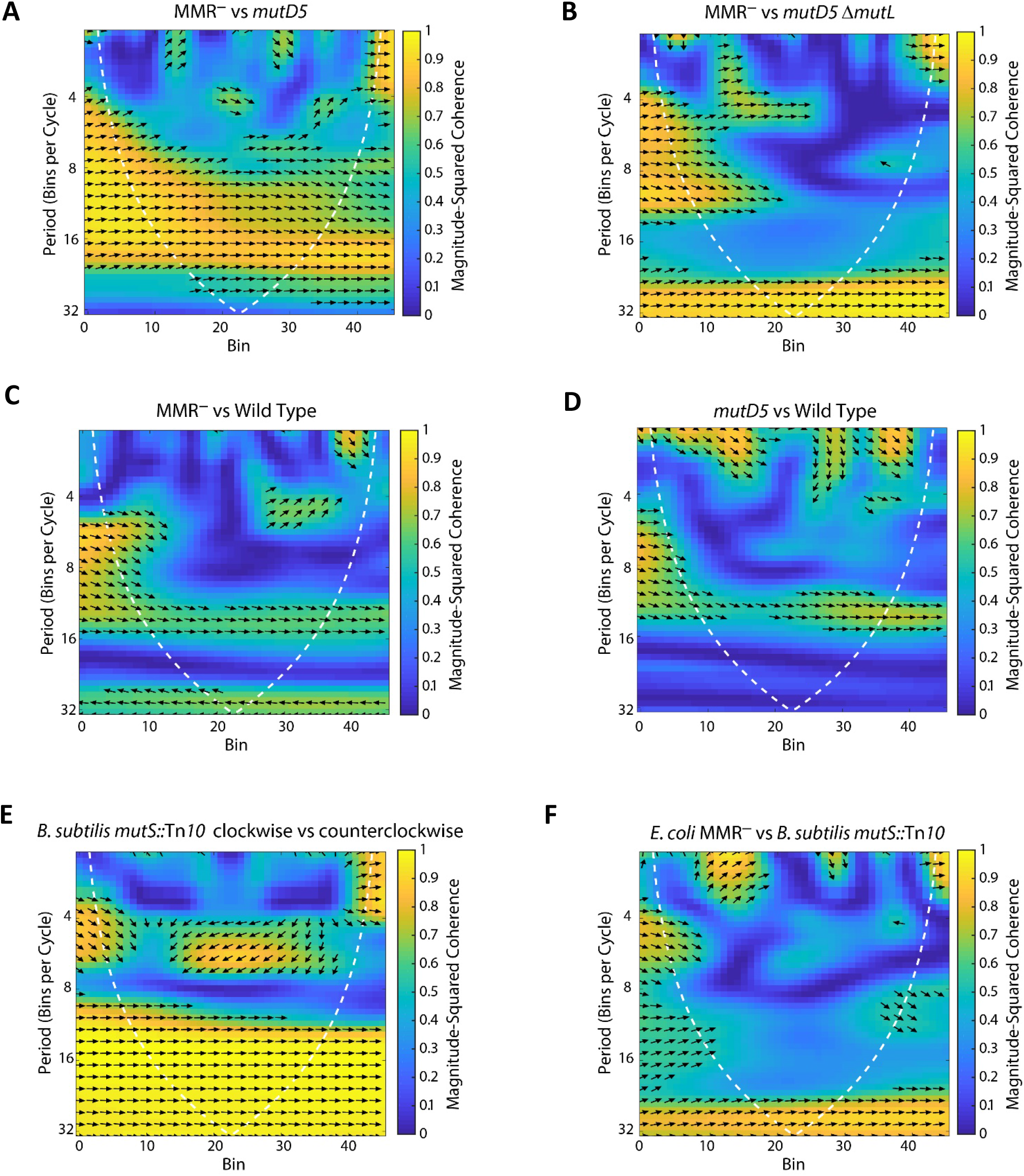
The BPS density pattern is a result of biased correction by replication proofreading and MMR. For a description of the plots, see the legend to Supplementary Figure S2. **Supplementary Figures S4A and B.** A strain with deficient proofreading but active MMR yields a BPS pattern similar to a MMR-defective strain, but a strain with neither proofreading nor MMR does not. The *mutD5* allele encodes an exonuclease-deficient proofreader. Strains: S4A, PFM163; S4B, PFM165/397/399. **Supplementary Figures S4C and D.** The BPS density pattern in the wild-type strain does not match the pattern of either the MMR-defective strain, or the mutD5 mutant strain. Strains: S4C and D, eight strains with wild-type mutational phenotypes (see text). **Supplementary Figures S4E and F.** MMR-defective *B. subtilis* also has a symmetrical BPS density pattern, but it is different than *E. coli’s* pattern. Strains: three *B. subtilis* mutant strains with the same mutational phenotype (see Supplemental Tables S1 and S3).

**Supplementary Figure S5.**
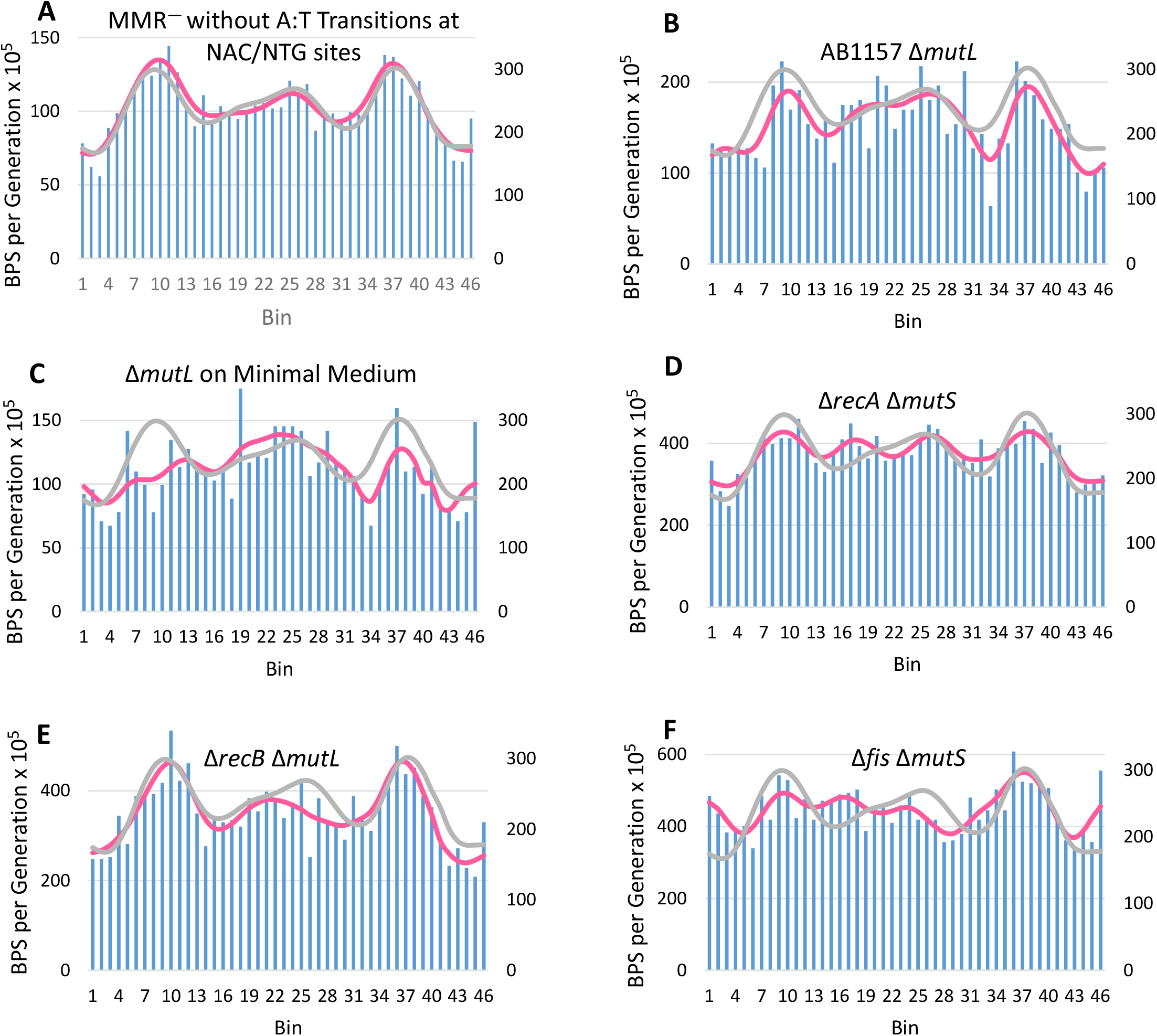
The BPS density patterns from experiments mentioned in the text but not shown. For each plot the data were collected into 100 Kb bins starting at the origin of replication on the left and continuing around the chromosome back to the origin. The BPS density patterns (bars) and Daubechies wavelet transforms (pink lines) of the indicated strains are compared to the Daubechies wavelet transform of the combined MMR^−^ strains (grey line). In each plot the left-hand scale has been adjusted to bring the wavelets close together for comparison. Strains: S5A, MMR strains but A:T transitions at 5’NAC3’/3’NTG5’ sites removed; S5B, PFM669; S5C, PFM5; S5D, PFM424; S5E, PFM456; S5F, PFM482.

**Supplementary Figure S6.**
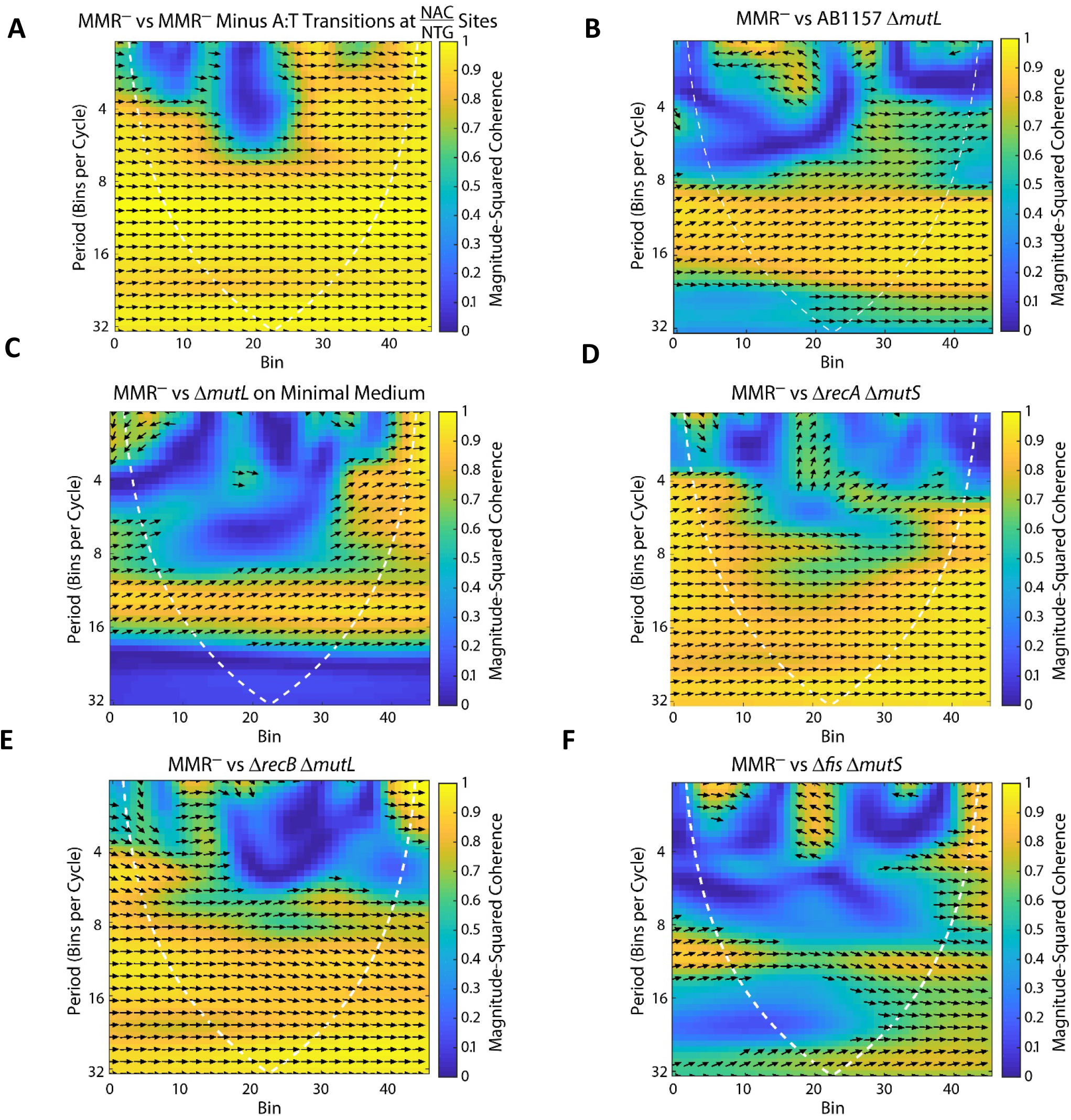
Plots of the wavelet coherence between the binned BPS data from the MMR-defective strains and the data from the experiments shown in Supplementary Figure S5. See the legends from Supplementary Figure S2 and S5 for details

## References

1. Foster PL, Hanson AJ, Lee H, Popodi EM, Tang H. On the mutational topology of the bacterial genome. G3 (Bethesda). 2013;3(3):399–407.

2. Dillon MM, Sung W, Lynch M, Cooper VS. Periodic variation of mutation rates in bacterial genomes associated with replication timing. MBio. 2018;9(4).

3. Dillon MM, Sung W, Sebra R, Lynch M, Cooper VS. Genome-wide biases in the rate and molecular spectrum of spontaneous mutations in *Vibrio cholerae* and *Vibrio fischeri*. Mol Biol Evol. 2017;34(1):93–109.

4. Wei W, Xiong L, Ye YN, Du MZ, Gao YZ, Zhang KY, et al. Mutation landscape of base substitutions, duplications, and deletions in the representative current cholera pandemic strain. Genome Biol Evol. 2018;10(8):2072–85.

5. Long H, Sung W, Miller SF, Ackerman MS, Doak TG, Lynch M. Mutation rate, spectrum, topology, and context-dependency in the DNA mismatch repair-deficient Pseudomonas fluorescens ATCC948. Genome Biol Evol. 2014;7(1):262–71.

6. Dettman JR, Sztepanacz JL, Kassen R. The properties of spontaneous mutations in the opportunistic pathogen Pseudomonas aeruginosa. BMC Genomics. 2016;17:27.

7. Lee H, Popodi E, Tang H, Foster PL. Rate and molecular spectrum of spontaneous mutations in the bacterium *Escherichia coli* as determined by whole-genome sequencing. Proceedings of the National Academy of Sciences of the United States of America. 2012;109(41):E2774–E83.

8. Marinus MG. DNA Mismatch Repair. EcoSal Plus. 2012; doi:10.1128/ecosalplus.7.2.5

9. Ganai RA, Johansson E. DNA replication-a matter of fidelity. Mol Cell. 2016;62(5):745–55.

10. Niccum BA, Lee H, MohammedIsmail W, Tang H, Foster PL. The spectrum of replication errors in the absence of error correction assayed across the whole genome of *Escherichia coli*. Genetics. 2018;209(4):1043–54.

11. Foster PL, Niccum BA, Popodi E, Townes JP, Lee H, MohammedIsmail W, et al. Determinants of base-pair substitution patterns revealed by whole-genome sequencing of DNA mismatch repair defective *Escherichia coli*. Genetics. 2018;209(4):1029–42.

12. Valens M, Penaud S, Rossignol M, Cornet F, Boccard F. Macrodomain organization of the *Escherichia coli* chromosome. EMBO J. 2004;23(21):4330–41.

13. Jinks-Robertson S, Bhagwat AS. Transcription-associated mutagenesis. Annu Rev Genet. 2014;48:341–59.

14. Rasouly A, Pani B, Nudler E. A magic spot in genome maintenance. Trends Genet. 2017;33(1):58–67.

15. Zhang X, Zhang X, Zhang X, Liao Y, Song L, Zhang Q, et al. Spatial vulnerabilities of the *Escherichia coli* genome to spontaneous mutations revealed with improved duplex sequencing. Genetics. 2018;210(2):547–58.

16. Withey JH, Friedman DI. A salvage pathway for protein structures: tmRNA and trans-translation. Annu Rev Microbiol. 2003;57:101–23.

17. Kogoma T. Stable DNA replication: interplay between DNA replication, homologous recombination, and transcription. Microbioland MolecBiolRev. 1997;61:212–38.

18. Maduike NZ, Tehranchi AK, Wang JD, Kreuzer KN. Replication of the *Escherichia coli* chromosome in RNase HI-deficient cells: multiple initiation regions and fork dynamics. Mol Microbiol. 2014;91(1):39–56.

19. Wang X, Lesterlin C, Reyes-Lamothe R, Ball G, Sherratt DJ. Replication and segregation of an *Escherichia coli* chromosome with two replication origins. Proceedings of the National Academy of Sciences of the United States of America. 2011;108(26):E243–50.

20. Ivanova D, Taylor T, Smith SL, Dimude JU, Upton AL, Mehrjouy MM, et al. Shaping the landscape of the *Escherichia coli* chromosome: replication-transcription encounters in cells with an ectopic replication origin. Nucleic Acids Res. 2015;43(16):7865–77.

21. Waldminghaus T, Skarstad K. The *Escherichia coli* SeqA protein. Plasmid. 2009;61(3):141–50.

22. Sutera VA, Jr., Lovett ST. The role of replication initiation control in promoting survival of replication fork damage. MolMicrobiol. 2006;60(1):229–39.

23. Sanchez-Romero MA, Busby SJ, Dyer NP, Ott S, Millard AD, Grainger DC. Dynamic distribution of SeqA protein across the chromosome of *Escherichia coli* K-12. MBio. 2010;1(1):e00012–10.

24. Weitao T, Nordstrom K, Dasgupta S. *Escherichia coli* cell cycle control genes affect chromosome superhelicity. EMBO Rep. 2000;1(6):494–9.

25. Lobner-Olesen A, Marinus MG, Hansen FG. Role of SeqA and Dam in *Escherichia coli* gene expression: a global/microarray analysis. Proceedings of the National Academy of Sciences of the United States of America. 2003;100(8):4672–7.

26. Waldminghaus T, Skarstad K. ChIP on Chip: surprising results are often artifacts. BMC Genomics. 2010;11:414.

27. Sun L, Fuchs JA. *Escherichia coli* ribonucleotide reductase expression is cell cycle regulated. Mol Biol Cell. 1992;3(10):1095–105.

28. Gon S, Camara JE, Klungsoyr HK, Crooke E, Skarstad K, Beckwith J. A novel regulatory mechanism couples deoxyribonucleotide synthesis and DNA replication in *Escherichia coli*. EMBO J. 2006;25(5):1137–47.

29. Cooper S, Helmstetter CE. Chromosome replication and the division cycle of *Escherichia coli* B/r. J Mol Biol. 1968;31(3):519–40.

30. Claret L, Rouviere-Yaniv J. Variation in HU composition during growth of *Escherichia coli:* the heterodimer is required for long term survival. J Mol Biol. 1997;273(1):93–104.

31. Dorman CJ. Function of nucleoid-associated proteins in chromosome structuring and transcriptional regulation. J Mol Microbiol Biotechnol. 2014;24(5-6):316–31.

32. Kahramanoglou C, Seshasayee AS, Prieto AI, Ibberson D, Schmidt S, Zimmermann J, et al. Direct and indirect effects of H-NS and Fis on global gene expression control in *Escherichia coli*. Nucleic Acids Res. 2011;39(6):2073–91.

33. Torrents E, Grinberg I, Gorovitz-Harris B, Lundstrom H, Borovok I, Aharonowitz Y, et al. NrdR controls differential expression of the *Escherichia coli* ribonucleotide reductase genes. J Bacteriol. 2007;189(14):5012–21.

34. Lane HE, Denhardt DT. The rep mutation. IV. Slower movement of replication forks in *Escherichia coli* rep strains. J Mol Biol. 1975;97(1):99–112.

35. Guy CP, Atkinson J, Gupta MK, Mahdi AA, Gwynn EJ, Rudolph CJ, et al. Rep provides a second motor at the replisome to promote duplication of protein-bound DNA. Mol Cell. 2009;36(4):654–66.

36. Boubakri H, de Septenville AL, Viguera E, Michel B. The helicases DinG, Rep and UvrD cooperate to promote replication across transcription units in vivo. EMBO J. 2010;29(1):145–57.

37. Myka KK, Hawkins M, Syeda AH, Gupta MK, Meharg C, Dillingham MS, et al. Inhibiting translation elongation can aid genome duplication in *Escherichia coli*. Nucleic Acids Res. 2017;45(5):2571–84.

38. Heller RC, Marians KJ. Non-replicative helicases at the replication fork. DNA Repair (Amst). 2007;6(7):945–52.

39. Sandler SJ, Marians KJ. Role of PriA in replication fork reativation in *Escherichia coli*. J Bact. 2000;182:9–13.

40. Duggin IG, Wake RG, Bell SD, Hill TM. The replication fork trap and termination of chromosome replication. MolMicrobiol. 2008;70(6):1323–33.

41. Mercier R, Petit MA, Schbath S, Robin S, El Karoui M, Boccard F, et al. The MatP/matS site-specific system organizes the terminus region of the *E. coli* chromosome into a macrodomain. Cell. 2008;135(3):475–85.

42. Espeli O, Borne R, Dupaigne P, Thiel A, Gigant E, Mercier R, et al. A MatP-divisome interaction coordinates chromosome segregation with cell division in *E*. coli. EMBO J. 2012;31(14):3198–211.

43. Syeda AH, Hawkins M, McGlynn P. Recombination and replication. Cold Spring Harb Perspect Biol. 2014;6(11):a016550.

44. Louarn JM, Louarn J, Francois V, Patte J. Analysis and possible role of hyperrecombination in the termination region of the *Escherichia coli* chromosome. J Bacteriol. 1991;173(16):5097–104.

45. Bierne H, Ehrlich SD, Michel B. The replication termination signal terB of the *Escherichia coli* chromosome is a deletion hot spot. EMBO J. 1991;10(9):2699–705.

46. Louarn J, Cornet F, Francois V, Patte J, Louarn JM. Hyperrecombination in the terminus region of the *Escherichia coli* chromosome: possible relation to nucleoid organization. J Bacteriol. 1994;176(24):7524–31.

47. Corre J, Cornet F, Patte J, Louarn JM. Unraveling a region-specific hyper-recombination phenomenon: genetic control and modalities of terminal recombination in *Escherichia coli*. Genetics. 1997;147(3):979–89.

48. Cox MM. Regulation of bacterial RecA protein function. Crit Rev Biochem Mol Biol. 2007;42(1):41–63.

49. Little JW, Harper JE. Identification of the lexA gene product of *Escherichia coli* K-12. Proceedings of the National Academy of Sciences of the United States of America. 1979;76(12):6147–51.

50. Prieto AI, Kahramanoglou C, Ali RM, Fraser GM, Seshasayee AS, Luscombe NM. Genomic analysis of DNA binding and gene regulation by homologous nucleoid-associated proteins IHF and HU in *Escherichia coli* K12. Nucleic Acids Res. 2012;40(8):3524–37.

51. Sobetzko P, Travers A, Muskhelishvili G. Gene order and chromosome dynamics coordinate spatiotemporal gene expression during the bacterial growth cycle. ProcNatlAcadSciUSA. 2012;109(2):E42–E50.

52. Cho BK, Knight EM, Barrett CL, Palsson BO. Genome-wide analysis of Fis binding in *Escherichia coli* indicates a causative role for A-/AT-tracts. Genome Res. 2008;18(6):900–10.

53. Wang W, Li GW, Chen C, Xie XS, Zhuang X. Chromosome organization by a nucleoid-associated protein in live bacteria. Science. 2011;333(6048):1445–9.

54. Lang B, Blot N, Bouffartigues E, Buckle M, Geertz M, Gualerzi CO, et al. High-affinity DNA binding sites for H-NS provide a molecular basis for selective silencing within proteobacterial genomes. Nucleic Acids Res. 2007;35(18):6330–7.

55. Bouffartigues E, Buckle M, Badaut C, Travers A, Rimsky S. H-NS cooperative binding to high-affinity sites in a regulatory element results in transcriptional silencing. Nat Struct Mol Biol. 2007;14(5):441–8.

56. Wolf SG, Frenkiel D, Arad T, Finkel SE, Kolter R, Minsky A. DNA protection by stress-induced biocrystallization. Nature. 1999;400(6739):83–5.

57. Foster PL, Lee H, Popodi E, Townes JP, Tang H. Determinants of spontaneous mutation in the bacterium *Escherichia coli* as revealed by whole-genome sequencing. Proceedings of the National Academy of Sciences of the United States of America. 2015;112(44):E5990–E599.

58. Duigou S, Boccard F. Long range chromosome organization in *Escherichia coli:* The position of the replication origin defines the non-structured regions and the Right and Left macrodomains. PLoS Genet. 2017;13(5):e1006758.

59. Dupaigne P, Tonthat NK, Espeli O, Whitfill T, Boccard F, Schumacher MA. Molecular basis for a protein-mediated DNA-bridging mechanism that functions in condensation of the *E. coli* chromosome. Mol Cell. 2012;48(4):560–71.

60. Lioy VS, Cournac A, Marbouty M, Duigou S, Mozziconacci J, Espeli O, et al. Multiscale structuring of the *E*. *coli* chromosome by nucleoid-associated and condensin proteins. Cell. 2018;172(4):771–83 e18.

61. Thiel A, Valens M, Vallet-Gely I, Espeli O, Boccard F. Long-range chromosome organization in *E. coli:* a site-specific system isolates the Ter macrodomain. PLoSGenet. 2012;8(4):e1002672.

62. Salgado H, Peralta-Gil M, Gama-Castro S, Santos-Zavaleta A, Muniz-Rascado L, Garcia-Sotelo JS, et al. RegulonDB v8.0: omics data sets, evolutionary conservation, regulatory phrases, cross-validated gold standards and more. Nucleic Acids Res. 2013;41(Database issue):D203–13.

63. Guo F, Adhya S. Spiral structure of *Escherichia coli* HUalphabeta provides foundation for DNA supercoiling. Proceedings of the National Academy of Sciences of the United States of America. 2007;104(11):4309–14.

64. van Noort J, Verbrugge S, Goosen N, Dekker C, Dame RT. Dual architectural roles of HU: formation of flexible hinges and rigid filaments. Proceedings of the National Academy of Sciences of the United States of America. 2004;101(18):6969–74.

65. Pinson V, Takahashi M, Rouviere-Yaniv J. Differential binding of the *Escherichia coli* HU, homodimeric forms and heterodimeric form to linear, gapped and cruciform DNA. J Mol Biol. 1999;287(3):485–97.

66. Bonnefoy E, Almeida A, Rouviere-Yaniv J. Lon-dependent regulation of the DNA binding protein HU in *Escherichia coli*. Proceedings of the National Academy of Sciences of the United States of America. 1989;86(20):7691–5.

67. Berger M, Farcas A, Geertz M, Zhelyazkova P, Brix K, Travers A, et al. Coordination of genomic structure and transcription by the main bacterial nucleoid-associated protein HU. EMBO Rep. 2010;11(1):59–64.

68. Browning DF, Grainger DC, Busby SJ. Effects of nucleoid-associated proteins on bacterial chromosome structure and gene expression. CurrOpinMicrobiol. 2010;13(6):773–80.

69. Grainger DC, Hurd D, Goldberg MD, Busby SJ. Association of nucleoid proteins with coding and non-coding segments of the *Escherichia coli* genome. Nucleic Acids Res. 2006;34(16):4642–52.

70. Blot N, Mavathur R, Geertz M, Travers A, Muskhelishvili G. Homeostatic regulation of supercoiling sensitivity coordinates transcription of the bacterial genome. EMBO Rep. 2006;7(7):710–5.

71. Benjamini Y, Hochberg Y. Controlling the false discovery rate-a practical and powerful approach to multiple testing. J Roy Stat Soc Ser B (Stat Method). 1995;57(1):289–300.

72. Grinsted A, Moore JC, Jevrejeva S. Application of the cross wavelet transform and wavelet coherence to geophysical time series. Nonlinear Processes in Geophysics. 2004;11:561–6.

73. Maraun D, Kurths J. Cross wavelet analysis: significance testing and pitfalls. Nonlinear Processes in Geophysics. 2004;11:505–14.

## Supplemental References

1. Lee H, Popodi E, Tang H, Foster PL. Rate and molecular spectrum of spontaneous mutations in the bacterium *Escherichia coli* as determined by whole-genome sequencing. Proceedings of the National Academy of Sciences of the United States of America. 2012;109(41):E2774–E83.

2. Dewitt SK, Adelberg EA. The Occurrence of a Genetic Transposition in a Strain of Escherichia Coli. Genetics. 1962;47(5):577–85.

3. Baba T, Ara T, Hasegawa M, Takai Y, Okumura Y, Baba M, et al. Construction of *Escherichia coli* K-12 in-frame, single-gene knockout mutants: the Keio collection. Mol Syst Biol. 2006;2:2006.

4. Miller JH. Experiments in molecular genetics. Cold Spring Harbor, N.Y.: Cold Spring Harbor Laboratory; 1972 1972.

5. Datsenko KA, Wanner BL. One-step inactivation of chromosomal genes in *Escherichia coli* K-12 using PCR products. Proc Natl Acad Sci USA. 2000;97:6640–5.

6. Miller JH. A short course in bacterial genetics: a laboratory manual and handbook for *Escherichia coli* and related bacteria. Cold Spring Harbor: Cold Spring Harbor Laboratory Press; 1992 1992.

7. Foster PL. Methods for determining spontaneous mutation rates. Methods Enzymol. 2006;409:195–213.

8. Hall BM, Ma CX, Liang P, Singh KK. Fluctuation analysis CalculatOR: a web tool for the determination of mutation rate using Luria-Delbruck fluctuation analysis. Bioinformatics. 2009;25(12):1564–5.

9. Foster PL, Lee H, Popodi E, Townes JP, Tang H. Determinants of spontaneous mutation in the bacterium *Escherichia coli* as revealed by whole-genome sequencing. Proceedings of the National Academy of Sciences of the United States of America. 2015;112(44):E5990–E599.

10. Foster PL, Niccum BA, Popodi E, Townes JP, Lee H, MohammedIsmail W, et al. Determinants of base-pair substitution patterns revealed by whole-genome sequencing of DNA mismatch repair defective *Escherichia coli*. Genetics. 2018;209(4):1029–42.

11. Li H, Durbin R. Fast and accurate short read alignment with Burrows-Wheeler transform. Bioinformatics. 2009;25(14):1754–60.

12. Wang X, Lesterlin C, Reyes-Lamothe R, Ball G, Sherratt DJ. Replication and segregation of an Escherichia coli chromosome with two replication origins. Proceedings of the National Academy of Sciences of the United States of America. 2011;108(26):E243–50.

13. Niccum BA, Lee H, MohammedIsmail W, Tang H, Foster PL. The spectrum of replication errors in the absence of error correction assayed across the whole genome of *Escherichia coli*. Genetics. 2018;209(4):1043–54.

14. Blank K, Hensel M, Gerlach RG. Rapid and highly efficient method for scarless mutagenesis within the *Salmonella enterica* chromosome. PLoS One. 2011;6(1):e15763.

